# Endogenous subthreshold noise and PIP_2_ tune diastolic coherence across the intact sinoatrial node through stochastic resonance

**DOI:** 10.64898/2026.09.18.752780

**Authors:** Manuel F. Muñoz, Marc D. Binder, Christopher G. Wilson, L. Fernando Santana

## Abstract

The heartbeat originates in the sinoatrial (SA) node, where spontaneously firing pacemaker myocytes interact to generate a coherent rhythm. The classical entrainment account is deterministic: the fastest cells entrain the rest, and variability is an error term that coupling suppresses. We proposed previously that the node instead exploits its own noise through stochastic resonance, with rhythm quality peaking at intermediate fluctuation amplitude, and since mapped a metabolic gradient capable of supplying it. Two-photon imaging of the voltage sensor ASAP5 in the intact mouse node resolved the subthreshold regime that tissue-scale mapping averages away. Diastolic coherence followed an inverted-U against endogenous noise amplitude in both poles, with optima 3.5-fold apart, and each pole operated near its own. β-adrenergic stimulation raised noise in both poles to a common level. The inferior pole, which began below its optimum, gained coherence, while the superior pole, already at its optimum, lost interval regularity. Blocking HCN-mediated I_f_ with ivabradine lowered superior coherence without changing superior noise, and lowered inferior noise without lowering inferior coherence, dissociating the two coordinates. Thus, coherence is a surface defined by noise and coupled-clock drive. Phosphoinositide 4,5-bisphosphate (PIP_2_), an ATP-dependent lipid cofactor for HCN channels, increased firing rate and noise in both poles but improved coherence only in the energy-poor inferior node. Prior I_f_ block abolished that rescue. Pacemaking is therefore graded by a bioenergetic supply chain that sets the noise, not by entrainment alone. Noise, long treated as a nuisance to be averaged away, is an active ingredient of the pacemaker.

**Key points summary:**

- The sinoatrial node contains pacemaker cells with different firing patterns and metabolic states, but how this heterogeneity produces a coherent heartbeat is not fully understood. Biological noise is usually viewed as something that disrupts coordination.
- Using two-photon voltage imaging in the intact mouse sinoatrial node, we found that subthreshold voltage fluctuations are structured, regionally graded and larger in the energy-poor inferior node.
- Coherence of SA node pacemaker cells was highest at an intermediate level of noise, demonstrating that endogenous noise can strengthen rhythm through stochastic resonance rather than simply impair pacemaking.
- Changing sympathetic drive, blocking the HCN pacemaker current and supplying phosphatidylinositol 4,5-bisphosphate (PIP_2_) showed that coherence depends jointly on noise amplitude and pacemaker drive, linking cellular energy supply to electrical coordination.
- These findings identify cardiac pacemaking as a bioenergetically tuned, noise-assisted process in which regional energy supply shapes subthreshold fluctuations and, together with pacemaker drive, determines coordination across the SA node.

## Introduction

The sinoatrial (SA) node, a crescent-shaped structure at the junction of the superior vena cava and right atrium in which clusters of myocytes expressing HCN (hyperpolarization-activated cyclic nucleotide-gated channels) generate spontaneous action potentials (APs), is the heart’s primary pacemaker. These APs propagate through gap junctions to neighboring SA node myocytes and ultimately to the working myocardium, setting heart rate. The spontaneous diastolic depolarization that drives SA node myocytes to threshold and initiates each cardiac cycle arises from the interplay of several conductances: the hyperpolarization-activated ‘funny’ current (I_f_) (DiFrancesco, 1986), carried by HCN4 channels (Fenske *et al*., 2020); L- and T-type Ca^2+^ currents, carried by Ca_V_1.2, Ca_V_1.3 and Ca_V_3.1; the Na⁺/Ca^2+^ exchanger (NCX) current, driven by rhythmic Ca^2+^ release from the sarcoplasmic reticulum via ryanodine receptors; and repolarizing K⁺ currents. These conductances generate a periodic impulse through a coupled system of membrane and Ca^2+^ clocks (Lakatta *et al*., 2010). Viewed at the level of its output, the node behaves as a near-metronome, with a single atrial and ventricular activation per cycle and a beat-to-beat interval variability (in the resting mouse) of a few percent. Tissue-level optical mapping largely preserves this impression of order, resolving activation as a wave that emerges from a leading site and spreads across the node along smooth isochrones (Efimov *et al*., 2010). How this process is perceived, however, depends on the resolution at which it is imaged.

When Ca^2+^ signals are resolved at the scale of individual cells, the smooth wave dissolves into a population of oscillators firing at different rates, interspersed with cells that fire irregularly or not at all, with local Ca^2+^ release distributed throughout the tissue (Bychkov *et al*., 2020; Bychkov *et al*., 2022). Isolated node myocytes show comparable electrical diversity, ranging from rhythmic firing to irregular firing and dormancy (Honjo *et al*., 1996; Kim *et al*., 2018; Grainger *et al*., 2021). Whether membrane potential behaves this way in individual cells *within* the intact node has been harder to establish because voltage in nodal tissue has conventionally been recorded either from single dissociated myocytes, which lose neighbor-to-neighbor contacts, or by optical mapping at a spatial scale that averages across many cells. Regardless of the answer, the dominant conceptual model for how such a population produces a single coherent rhythm is entrainment: the fastest cells, acting as the leading pacemaker, phase-lock slower neighbors through electrotonic coupling (Jalife, 1984; Anumonwo *et al*., 1991). In this view, the order visible at low resolution is the signal, and the variability visible at high resolution is degradation to be minimized.

The heterogeneity of the node motivates a re-examination of this model, because the model oversimplifies the complex, multi-layered mechanisms that drive pacemaker tissue. Anatomical and molecular mapping have established graded differences in myocyte size, ion channel and connexin expression, and intercellular coupling between the nodal center and periphery, giving rise to the classical ‘gradient’ and ‘mosaic’ formulations of nodal organization (Honjo *et al*., 1996; Dobrzynski *et al*., 2005; Tellez *et al*., 2006). High-resolution optical mapping in canine, human, and rodent hearts has subsequently shown that this structural heterogeneity is functionally partitioned, demonstrating that the node contains multiple intranodal pacemaker sites and discrete SA conduction pathways whose engagement shifts with autonomic tone, ionic conditions and disease, conferring redundancy that protects the node from failure (Fedorov *et al*., 2009; Fedorov *et al*., 2010; Glukhov *et al*., 2010; Csepe *et al*., 2016; Li *et al*., 2017). In rodents, this partitioning follows a superior–inferior axis: both regions are capable of pacemaking, but the superior region is normally dominant, and the inferior region assumes leadership only when the superior node is suppressed (Brennan *et al*., 2020). Quantitative proteomic and single-nucleus transcriptomic surveys have since shown that nodal myocytes are a molecularly diverse rather than a uniform population (Linscheid *et al*., 2019), and cell-resolved Ca^2+^ imaging has revealed that a substantial fraction of nodal cells are dormant or fire irregularly, with subthreshold Ca^2+^ fluctuations distributed throughout the tissue (Bychkov *et al*., 2020; Bychkov *et al*., 2022). These are features that a purely deterministic entrainment framework accommodates awkwardly.

Superimposed on these anatomical, molecular, and electrical layers is an energetic layer. The transition from superior to inferior node is accompanied by a decline in microvascular density and mitochondrial content and an increase in myocyte-to-vessel distance; as a result, superior myocytes generate larger, faster, beat-locked ATP transients, while inferior myocytes operate in a lower-gain or energy-deficit mode (Grainger *et al*., 2021; Manning *et al*., 2025; Muñoz *et al*., 2026). Together, these features define a capillary → mitochondrion → ion channel supply chain whose capacity varies with position in the node (Santana & Earley, 2026). How this gradient impacts the dynamics of the rhythm, as opposed to the mean firing rate, has not been addressed. The distinction matters because fluctuation behavior and mean firing rate vary independently across the node (Grainger *et al*., 2021), indicating that these are separable properties rather than two readouts of the same gradient.

Here, we test the hypothesis that the same gradient sets the level of subthreshold membrane noise, and that rather than being an incidental problem, this noise, in the form of stochastic resonance, is functionally important. Stochastic resonance arises when an intermediate level of random noise improves rather than degrades the detection or generation of a periodic signal. A system that meets three conditions—a threshold, a weak periodic drive, and a tunable source of noise—can exhibit stochastic resonance (Bezrukov & Vodyanoy, 1995; Clancy & Santana, 2020; Guarina *et al*., 2022; Pérez-Cervera *et al*., 2023; Kreider *et al*., 2025). The diastolic depolarization of a pacemaker cell approaching its firing threshold satisfies all three. The firing threshold provides the nonlinearity; the rhythmic subthreshold Ca^2+^ releases of the Ca^2+^ clock, acting through NCX, provide the weak periodic drive to the membrane; and stochastic gating of the conductances active during diastole provides the noise. Identifying the drive in this way distinguishes the phenomenon from coherence resonance, in which a noise optimum arises in an autonomous oscillator with no periodic input at all (Pikovsky & Kurths, 1997).

Recent single-cell and *in silico* work by Okamura *et al*. (2026) established this principle, demonstrating that white-noise currents injected into isolated rabbit SA node myocytes optimize firing regularity along an inverted-U relationship and prevent modeled tissue-level sinus arrest. Three open questions remain: 1) Because the functional experiments reported by Okamura *et al*. (2026) delivered noise through an electrode, whether endogenous fluctuation, which is neither white nor externally timed, imposes the same regularity was not tested. 2) The tissue-level claim rests on simulation, and the accompanying intact-tissue imaging characterized the Ca^2+^ noise environment rather than asking whether resonance organizes rhythm among cells. 3) Finally, cells were sorted by firing behavior rather than anatomical origin, leaving the question of whether a cell’s position on the resonance curve is set by its local energetic state unaddressed.

Here, we address all three of these open questions. Using two-photon imaging, we resolved membrane potential in individual myocytes across the intact node, bringing into view the subthreshold regime that tissue-scale mapping averages away. Diastolic coherence is maximized at an intermediate level of endogenous subthreshold noise, with the superior and inferior poles each sitting near their own optimum. A pole’s position on this resonance curve is dictated by its local energy supply. We identified phosphoinositide 4,5-bisphosphate (PIP_2_) as the mechanistic link, showing that its ATP-dependent synthesis by phosphatidylinositol 4-kinase (PI4K) constitutes a metabolic sensor positioned directly in the pathway setting pacemaker drive. Supplying exogenous PIP_2_ accelerated firing globally but relieved the coherence limitation only in the ATP-poor inferior node. The SA node is therefore not a population of oscillators whose variability degrades a deterministic rhythm. It is a spatially graded structure where noise is shaped by metabolic capacity and tuned to sharpen the beat.

## Materials and Methods

### Ethical approval

All animal procedures were approved by the Institutional Animal Care and Use Committee (IACUC) of the University of California, Davis (protocol #24065) and were conducted in accordance with the National Institutes of Health (NIH) *Guide for the Care and Use of Laboratory Animals*.

### Animals and viral expression

Male, wild-type, 8–12-wk-old C57BL/6J mice (The Jackson Laboratory, USA) were used for all experiments in this study. Pacemaker-cell–specific expression of the genetically encoded voltage indicator ASAP5 (Hao *et al*., 2024) was achieved by infecting mice with an AAV9 vector in which ASAP5 expression is driven by the *HCN4* promoter, restricting the indicator to HCN4^+^ SA node myocytes. Mice were euthanized with a single intraperitoneal administration of a lethal dose of sodium pentobarbital (250 mg/kg).

### Two-photon line-scan imaging

*Ex vivo* SA node preparations were imaged on an inverted two-photon microscope as previously described in detail (Muñoz *et al*., 2026). Briefly, ASAP5 was excited at 850 nm and imaged using an Olympus FV3000 MPE-RS multi-photon microscope equipped with an XL Plan N 25X lens (NA = 1.05) in line-scan mode (2 ms/line with a pixel size of 1 µm). During acquisition, preparations were continuously perfused *ex vivo* at room temperature with a Tyrode III solution supplemented with 10 μM blebbistatin to suppress contractile motion. For pharmacological experiments, ivabradine (50 μM; Tocris, USA), isoproterenol (100 nM; Sigma, USA), or diC8-PIP_2_ (10 μM; Sigma) was added directly to the perfusate.

### GEVI LineScan Analyzer software

All line-scan images were analyzed with the custom, browser-based application, GEVI LineScan Analyzer (v1.8.1), developed in this study. Portions of the source code were drafted with the assistance of a large language model (Claude Code, Anthropic). All code was reviewed, modified, tested, and validated by the authors, who take responsibility for its correctness. All decisions regarding analytical approach, parameter selection, and interpretation of the data were made by the authors. No generative artificial-intelligence tool was used to produce, alter, or interpret the experimental data, and no such tool meets the criteria for authorship. The application is a single self-contained HTML/JavaScript file that runs entirely client-side in any modern web browser, with no installation, server, or upload of data required; recordings are processed locally in the browser. The application integrates the complete workflow: image loading and acquisition calibration (ms/line, µm/pixel), sensor selection and F/F_0_ conversion, photobleach correction, event detection, AP/subthreshold classification, and the full entrainment/stochastic-resonance analysis suite (Kuramoto coherence, subthreshold-noise σ, power spectral density, spatial autocorrelation, conditional firing probability, phase-timing, shuffled controls, noise–regularity mapping, leading-site stability, conduction velocity, and entropy), and exports all per-recording results as a single JSON object and as CSV tables for downstream aggregation. Every setting that governs an analysis is written to the parameters block of that JSON export, including analyzer version, sensor, ms/line and µm/pixel and their source, photobleach model and its parameters, detection z-score threshold, AP-threshold mode and value, and whether per-window detrending was enabled, so every value reported here can be traced to the settings that produced it. To support reuse and reproducibility, we are releasing the Analyzer as open-source software rather than as Supplemental Software; source code for v1.8.1 and all subsequent versions is maintained in a public repository under the GNU General Public License v3.0 (copyright, The Regents of the University of California), and the version used for every analysis reported here is archived under a persistent DOI (see *Data and Software Availability*).

### Signal extraction and event detection

For the ASAP sensor, fluorescence was corrected for photobleaching with an exponential decay model and then converted to ΔF/F_0_. F_0_ was determined separately for each spatial column as the lowest fluorescence value reached during diastole within an epoch of at least 10 ms over which fluorescence was flat. One F_0_ was assigned per column and held fixed for the whole recording. No time-varying or per-cycle baseline entered the normalization. Voltage events were detected per spatial column using a dual-polarity SparkMaster 2 detection engine (Tomek *et al*., 2023) applying the following multi-step pipeline adapted for voltage-indicator line scans: (1) optional extracellular masking, (2) column normalization, (3) row normalization (drift removal), (4) spatial filtering (2D median + 1D Gaussian), (5) an SD-transform producing a per-pixel z-score image, and subsequent connected-component labeling, amplitude/duration gating, and event formatting. The engine runs two symmetric passes (positive- and negative-going z) to capture both polarities without sensor-specific threshold tuning, using a z-score detection floor combined with a compound-score criterion and an event-duration filter (APs typically ≥ 80 ms; subthreshold events ≥ 20 ms). The engine is a derived work of SparkMaster 2, reimplemented in JavaScript and retuned for voltage-indicator data. It is redistributed as part of the Analyzer under the GNU General Public License v3.0, and all analyses downstream of detection are original to this study.

### Detection validation

Detection performance was measured against seeded synthetic line-scan images containing planted events of known time, amplitude, and duration. Over a defined operating range optimized for ASAP5 SA node recordings (AP amplitude ≥ 0.45 ΔF/F_0_; signal-to-noise ratio [SNR] ≥ 8), the detection engine achieved 100% sensitivity and 98.4% precision (1.6% false-positive rate). Default parameters were set to this operating point and adjusted only when imaging conditions deviated from it. The synthetic line-scan generator (a phase-delayed, multi-column beating pattern with additive noise) is built into the Analyzer and exercises the full pipeline from detection through entrainment analysis. It is seeded, so the validation reported here can be reproduced from the archived software release without access to the experimental recordings. AP-versus-subthreshold classification was derived from the bimodal amplitude histogram using automatic valley detection, with a depth-validation safeguard (the valley must fall below 60% of the shorter peak) and optional manual or population-fixed threshold modes; the method used for each recording was logged in the export metadata.

### Amplitude classification (APs vs. subthreshold fluctuations)

Detected events were classified into two amplitude populations from the bimodal ΔF/F₀ histogram. The boundary between the leftmost (subthreshold) and rightmost (AP) peaks was placed at the histogram valley (automatic valley detection, with manual or population-fixed options). Events above the boundary were labeled APs; all remaining detected events were labeled SVFs. No independent lower amplitude floor was imposed on the subthreshold class.

### Firing-mode classification (periodic, irregular, non-firing)

Beat-to-beat regularity was quantified with an irregularity score (IS) computed from the AP interval series of each line-scan recording. For successive inter-AP intervals, ISI_n_, the score for each interval pair was IS_n_ = 100 × |ISI_n_ − ISI_(n−1)_|/ISI_(n−1)_, and each cell was summarized by the median IS across its intervals (Telgkamp *et al*., 2002; Zanella *et al*., 2014). A local, difference-based score was preferred over the CV because it is insensitive to slow drift in mean firing rate within a recording.

Cells with fewer than three detected APs were classified as non-firing; cells with fewer than four intervals were not classified. For the remaining cells, the periodic/irregular boundary was derived from the data rather than assumed by fitting a two-component Gaussian mixture to the distribution of log_10_(median IS) pooled across all cells and both poles and placing the threshold at the crossing point of the two fitted components. This yielded IS = 24.0, corresponding to an interval CV of ∼0.25 by an IS-to-CV calibration. Cells with a median IS below this threshold were classified as periodic and those at or above it, as irregular. Sensitivity to this choice was assessed by sweeping the threshold and recomputing the proportion of periodic cells in each pole (**Figure 2B, C**).

### Diastolic coherence (Kuramoto order parameter)

For each time point t, every valid column was assigned a phase from its position within the current inter-AP interval (time since previous AP relative to the interval to the next AP), giving the phase variable required for the Kuramoto order parameter. Columns are line-scan positions rather than segmented cells, so adjacent columns sampling a single cell are not independent; therefore, R is computed across columns rather than across identified cells. The spatial autocorrelation length λ (**Figure 3B**) sets the distance over which neighboring columns carry shared signal and so bounds the effective number of independent units contributing to R. Phase is defined by supra-threshold APs only. The bracketing events used to compute φ are the AP peaks that satisfy the amplitude and duration criteria of the detection engine, so SVFs do not enter the phase calculation; they do contribute to noise measures (σ, PSD, entropy) but not to R. A column contributes to R(t) only when t falls between two detected APs of that column. Columns that are silent, or whose current interval is not bracketed by two APs (e.g., a missed or skipped beat) have no defined phase at t and are excluded from the average at that instant rather than assigned an arbitrary value. The instantaneous order parameter R(t) = |⟨*e*^iφx(t)^⟩| was averaged over the contributing columns, and the mean diastolic R was computed over diastolic epochs (excluding the AP upstroke). R = 1 indicates perfect phase-locking; R = 0 indicates fully scattered phases. Because φ is defined as fractional position within each column’s current interval, R is insensitive to interval irregularity shared across columns: a set of columns following one identically irregular AP train occupies the same phase at every instant and yields R = 1. R declines when columns disagree with one another, whether through independent beat-to-beat jitter, a systematic activation lag between columns, or heterogeneity in firing rate across columns.

Because a fixed activation lag lowers R independently of beat-to-beat jitter, conduction delay across the imaged field could in principle reduce R without any loss of coordination. R was therefore also computed on lag-corrected timings. For each column, the mean activation time relative to the earliest-activating column of the same recording was estimated across all beats and subtracted as a fixed offset, and phases were recomputed from the residual timings, so that the corrected R reflects timing dispersion about the mean activation sequence rather than the sequence itself. R is reported with and without this correction, and the spread of the mean activation profile across columns is reported by pole. Interval regularity is reported separately, indexed by CV-IBI (**Supplementary Figure 1**).

### Subthreshold noise, spectra, and spatial structure

Subthreshold noise amplitude was quantified as σ, the standard deviation of the diastolic voltage signal about its own local trend. Because both the diastolic depolarization ramp and the duration of diastole vary with firing rate, and both inflate an SD computed over whole diastoles, σ was measured in a fixed-length window placed at a fixed phase of every diastole. For each diastolic interval, defined from the end of one AP to the onset of the next, a window of fixed duration (100 ms by default) was placed to end a fixed offset (20 ms) before the next AP onset; diastoles shorter than the window plus offset were excluded. Within each window, a straight line was fitted by least squares and subtracted, and σ was taken as the standard deviation of the pooled residuals for that spatial column, then averaged across columns within the ROI. This measure is therefore both ramp-subtracted and rate-controlled: it reports the residual fluctuation about the deterministic diastolic trajectory, over a window of identical length and identical phase in every cell, and is independent of cycle length. The number of qualifying windows per column and per recording is reported with the source data. σ is a property of the continuous subthreshold trace and is distinct from the amplitude of the discrete subthreshold events classified above; the two are reported separately and are not interchangeable.

The power spectral density of the subthreshold signal was estimated and the aperiodic exponent β extracted by fitting P(f) ∝ f^−β^. Because firing rate shifts the beat fundamental and its harmonics through the fitting band, β was estimated over a rate-matched band identical for both poles, defined as a fixed multiple of the beat frequency, with the fundamental and its first harmonics notched out by spectral parameterization so that oscillatory peaks do not enter the aperiodic fit. Reference values: β ≈ 0 corresponds to white noise, β = 1 to pink noise, and β = 2 to Brownian noise. Spatial autocorrelation of the subthreshold signal yielded a spatial correlation length λ, defined as the separation at which the autocorrelation falls to 1/e. Because the myocyte long-axis orientation relative to the scan line was not controlled, λ is a mixed within- and between-cell measure and is interpreted as a relative rather than an absolute length; poles were compared under identical acquisition and analysis conventions.

For the regional comparisons in **Figure 3**, σ was computed as described above. The same regional difference was present in the uncorrected, non-detrended whole-diastole measure, provided as a supplementary quantification. For the optical and motion controls, intact preparations expressing a HaloTag-containing reporter were incubated for 10 min with 0.1 µM JFX554 HaloTag ligand (#HT1030; Promega, USA). JFX554-HaloTag fluorescence was acquired in line-scan mode from fixed regions of interest, converted to baseline-normalized traces, and passed through the pipeline used for subthreshold ASAP5 to yield σ, the aperiodic spectral exponent β, and permutation entropy in superior and inferior regions.

### Rate independence of σ and β

Because both σ and the spectral exponent β can be biased by firing rate, all regional and noise– coherence comparisons that use them were computed under the rate-controlled conventions defined above. A subthreshold signal accumulates variance in proportion to the length of the window over which it is measured, at a rate set by β, so the longer diastoles of slow-firing cells would inflate an SD-based σ independently of any true amplitude difference. At the measured exponent of β ≈ 1.5, σ scales as T^0.25^, so a diastolic interval ∼3-fold longer inflates σ by about 30%, against a measured regional σ ratio of ∼2.6-fold. The fixed-window, ramp-subtracted definition removes this bias by construction, and the regional gradient is present in the uncorrected measure as well.

The σ–coherence relationship was additionally tested in a linear mixed-effects model of diastolic R on σ and σ^2^, with firing rate and diastolic duration as fixed covariates and recording (or animal) as a random effect, and was examined within narrow rate bins and within each pole separately as within-pole controls. Because the poles differ in drive, coupling and diastolic input impedance, and therefore in the position of their optima, curves were fitted separately for each pole rather than pooled across poles.

### Stochastic-resonance (noise–regularity) mapping

Subthreshold noise amplitude and rhythm quality were paired at the level of the recording, with each recording contributing one mean σ and one diastolic R (and one CV-IBI). The relationship was fitted with a quadratic (R = a·σ^2^ + b·σ + c) by least squares separately within each anatomical pole; a negative curvature (a < 0) defines an inverted-U whose vertex gives the coherence-optimizing noise level for that pole. Curvature was assessed from the coefficient on the quadratic term and its 95% CI, and vertices are reported. Poles were not pooled, since their optima are region-specific, and the equivalent fit to CV-IBI defines the corresponding U, shown in **Supplementary Figure 1**.

### Rhythm entropy and noise-complexity measures

Entropy measures were computed in the GEVI LineScan Analyzer v1.8.1, separating timing irregularity from subthreshold-noise complexity. The analyzer provides three such measures: (i) ISI Shannon entropy, determined for each spatial column by binning (12 bins) the inter-spike-interval (ISI) series of detected APs and computing its Shannon entropy, H = −Σ p_i_ log_2_ p_i_, in bits. A global ISI entropy was also computed by pooling ISIs across columns, and the ISI coefficient of variation (CV = SD/mean) was reported per column as a complementary irregularity index. (ii) Permutation entropy (Bandt–Pompe; order m = 3, delay = 1; normalized 0–1) of the diastolic (non-AP) subthreshold ΔF/F_0_ trace, an ordinal-complexity measure of the subthreshold noise itself. (iii) Sample entropy (m = 2, r = 0.2·SD) of the spatially-averaged subthreshold trace, as a regularity measure of the noise. Of these, only permutation entropy data is presented in the paper. The ISI Shannon entropy/CV and sample entropy outputs are described here for completeness but are not shown. Per-recording means of permutation entropy were plotted against subthreshold noise amplitude (σ) to test for a U-shaped (entropy-minimizing) relationship and compared between superior and inferior node.

### Electrical field stimulation and voltage imaging of the isolated SA node

Electrically evoked activity was induced in isolated SA node preparations by field stimulation using two platinum wire electrodes positioned on opposite sides of the tissue within the perfusion chamber. SA nodes were prepared as described above and continuously superfused with Tyrode solution containing 10 µM blebbistatin to limit motion. ASAP5 fluorescence was first recorded during spontaneous sinus rhythm and following sequential treatment with ivabradine and diC8-PIP_2_. After pharmacological treatment, voltage pulses (10 V, 6 ms duration) were delivered at 3 Hz using a Grass stimulator (AstroMed Inc., USA) to confirm that the SA node tissue remained electrically excitable and capable of generating evoked APs.

### Optical mapping

After confirmation of deep anesthesia, hearts were rapidly isolated and transferred to warmed Tyrode solution consisting of 130 mM NaCl, 5.4 mM KCl, 1.8 mM CaCl_2_, 0.33 mM NaH_2_PO_4_, 25 mM HEPES, 22 mM glucose, and 0.5 mM MgCl_2_, with pH adjusted to 7.4 using NaOH. The SA node was then dissected from the heart and pinned flat for optical recordings. Membrane potential changes were assessed by first loading SA node preparations with the voltage-sensitive dye RH237 (10 µM; Thermo Fisher Scientific) for 20 min at 37°C and washing for 10 min in dye-free Tyrode solution containing 10 µM blebbistatin (Cayman Chemical) to prevent motion artifacts. The SA node preparation was then placed in the optical mapping setup (MappingLab, UK) and continuously perfused at 2.5 mL/min with oxygenated Tyrode solution, maintained at 37°C and equilibrated with a gas mixture of 95% O_2_ and 5% CO_2_. Optical mapping recordings in a 100 × 100 pixel acquisition region were obtained using a high-speed sCMOS camera (Prime BSI Express, Teledyne Photometrics). For voltage imaging, SA node preparations loaded with RH237 were excited through a 530/20 nm excitation filter. For measurement of membrane voltage changes, emitted fluorescence was separated using a 638 nm dichroic mirror and passed through a 700-nm long-pass emission filter. Fluorescence images were acquired at a sampling rate of 1000 frames/s. Optical mapping recordings were acquired and analyzed using OMapScope 5 software (version 5.9.41; MappingLab).

### Additional analyses

Leading-pacemaker-site stability, apparent conduction velocity (leading-edge analysis; valid cells: conduction velocity ≥ 10 µm/ms, cell width ≤ 200 µm), conditional firing probability, and phase-timing of subthreshold fluctuations were computed as implemented in the GEVI LineScan Analyzer v1.8.1.

### Statistical Analysis

Data were analyzed using two statistical frameworks according to the experimental design. For the baseline survey (**Figures 2** to **4**), a single recording from either the superior or inferior SA node was obtained from each animal; observations were therefore independent, with no random effect to estimate. Regional comparisons in this cohort were analyzed using unpaired two-sample tests. The coherence-noise quadratics of **Figure 4D** to **4F** were fitted by ordinary least squares separately within each pole, with every observation weighted equally and treated as independent, and with firing rate added as a fixed covariate to confirm that the curvature was not a rate effect.

For the intervention experiments (**Figures 5** to **9**), repeated measurements were obtained from the same preparation across treatment conditions. **Figures 5** to **8** involved two conditions, whereas **Figure 9** used a sequential three-condition protocol consisting of baseline, ivabradine, and ivabradine + diC8-PIP_2_. Thus, observations were paired within preparation. Within-pole drug effects were assessed by paired t-tests with Šídák adjustment for the two tests performed on each readout, and the pole × drug interaction was fitted as a linear mixed-effects model with preparation as a random effect, which accounts for the correlation between repeated measurements made on the same preparation and prevents pseudo-replication.

Firing rate and diastolic duration were included as fixed covariates where noted. The σ–coherence relationship was additionally examined within narrow firing-rate bins as a within-pole control. Curvature was tested by the coefficient on the quadratic term rather than by a model-level F-test, since the model F is significant for a purely monotonic relationship. To ensure regional noise differences were not artifacts of longer diastolic intervals, we included firing rate and diastolic duration as covariates wherever the design permitted their estimation. Exact *P-*values, F-statistics, degrees of freedom, and 95% CIs are reported directly in the main text and figure legends. Statistical significance was defined as *P* < 0.05.

## Results

### Voltage imaging resolves two classes of electrical events in single cells of the intact node

Much of what we know about the biophysical mechanisms of cardiac pacemaking comes from cells isolated from the node and studied individually. To visualize pacemaking as it unfolds, within the intact SA node and in many neighboring cells at once, we expressed the genetically encoded voltage indicator ASAP5 (Accelerated Sensor of Action Potentials) under control of the HCN4 promoter, delivered by an AAV9 vector, which restricts the indicator to HCN4-expressing SA node pacemaker myocytes. ASAP brightens and dims as the cell’s membrane voltage increases and decreases, providing a direct optical readout of the cell’s electrical activity in the form of a fluorescence trace. All experiments in this study were performed in the presence of 10 µM blebbistatin to arrest contraction and prevent motion artifacts, as is the standard in optical mapping of cardiac tissue (Fedorov *et al*., 2010; Ferreiro *et al*., 2012; Swift *et al*., 2021; Muñoz *et al*., 2026).

*Ex vivo* preparations of the intact node were imaged on an inverted 2-photon microscope in line-scan mode, in which the microscope repeatedly sweeps a single line across the tissue (one line every 2 ms at 1 µm per pixel) rather than building a full two-dimensional image. Trading the second spatial dimension for speed allows us to follow voltage in each cell along that line with millisecond timing—fast enough to resolve individual heartbeat cycles simultaneously in every cell (**Figure 1A–C**). This is what distinguishes our measurement from conventional optical mapping of the node, which averages fluorescence over regions containing hundreds of cells: because subthreshold fluctuations differ in timing from cell to cell, spatial averaging cancels them and leaves only the synchronous AP. Resolving events below threshold requires reading each cell separately, which is why the subthreshold signal described here has not previously been accessible in intact tissue.

**Figure 1.**
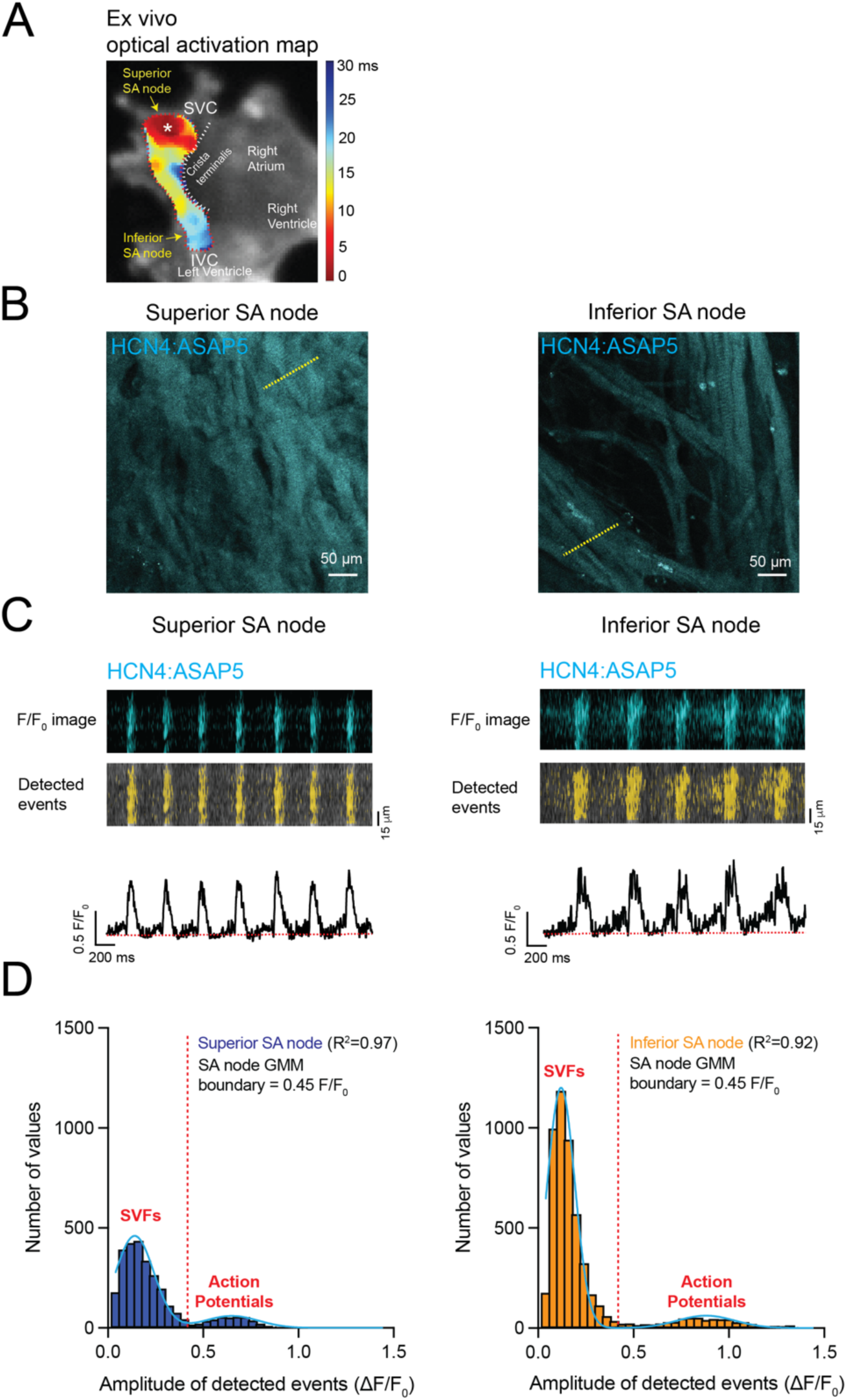
Two-photon ASAP imaging resolves two voltage-event populations in the intact SA node. (A) *Ex vivo* optical activation map of the whole node showing a smooth superior → inferior wave (superior and inferior SA node, SVC/IVC, crista terminalis marked). (B) HCN4:ASAP5 two-photon fields for the superior and inferior node with the line-scan position (dashed line). (C) Representative line-scan F/F_0_ image, masked image, and extracted ΔF/F_0_ trace for each pole. (D) Bimodal ΔF/F_0_ amplitude histograms fitted by a two-Gaussian mixture (superior R^2^ = 0.97; inferior R^2^ = 0.92) with the AP/SVF boundary at 0.45 ΔF/F_0_; SVFs (left) and APs (right).

At low magnification, optical-activation maps of the whole node showed a smooth wave of depolarization sweeping from the superior toward the inferior pole. This is the textbook picture of an orderly pacemaker (**Figure 1A**). At single-cell resolution, however, the same activity was strikingly noisy: between full APs, cells produced frequent small depolarizing events (**Figure 1C**). This raises the first question the paper addresses: are those small events meaningful, or just measurement jitter?

As a first step to addressing this, we detected and measured events with the GEVI LineScan Analyzer v1.8.1, an open-source browser application developed for this work and released with this article (see Materials and Methods). Its detection algorithm is adapted from SparkMaster 2 (Tomek *et al*., 2023), built for the analogous problem of identifying small, stochastic, spatially localized Ca^2+^-release events against a noisy line-scan background. Pooling all detected events, we found that ΔF/F_0_ amplitudes formed a clearly bimodal distribution, two peaks rather than one that were well fit by a two-component Gaussian mixture (superior R^2^ = 0.97; inferior R^2^ = 0.92) (**Figure 1D**). The two peaks correspond to a small population of large events—the full APs that drive the beat—and a much larger population of small events we term subthreshold voltage fluctuations (SVFs). A single amplitude boundary at 0.45 ΔF/F_0_ separated them; there is no distinct lower floor, so SVFs are simply every detected voltage event below the AP threshold. Both poles produced both event classes, but their balance differed, with the inferior node generating proportionally more SVFs relative to APs.

Pacemaker cells in the intact SA node therefore operate with two well-separated classes of voltage event: beats and (a much larger number of) small subthreshold fluctuations, with the ratio of the two already hinting at a regional difference. These discrete events constitute one component of a broader subthreshold signal: the diastolic trace between beats also carries slower, continuous fluctuations that no threshold-based classification resolves. In the remainder of the paper, we treat the entire subthreshold signal—SVFs and background together—as a candidate variable in setting the rhythm.

### Firing shifts from periodic to irregular along the superior–inferior axis

Having established that cells produce beats plus SVFs, we asked whether the regularity of the beats varies with position in the node. Functional heterogeneity along the superior–inferior axis is well documented and is not in itself contentious (Bychkov *et al*., 2020; Maltsev & Stern, 2022). Pacemaker myocytes isolated from the two poles differ in firing rate and regularity under current clamp, a recording in which both APs and subthreshold depolarizations are resolved directly (Grainger *et al*., 2021). What has not been established is whether this electrical heterogeneity survives in the intact, electrically coupled node, where mutual entrainment might be expected to dissolve it into a common rhythm, and whether it is graded systematically with position rather than distributed randomly among cells. Cell-resolved imaging of the whole mount allows us to ask both questions directly.

Representative line scans and corresponding voltage traces revealed distinct patterns of spontaneous activity across the two poles (**Figure 2A**). Cells in the superior SA node commonly displayed periodic AP firing, with successive APs occurring at relatively uniform intervals. In contrast, cells in the inferior SA node more frequently showed irregular AP firing, characterized by marked beat-to-beat variation in the timing of successive APs. A smaller subset of cells in both regions remained non-firing during the recording period, displaying SVFs without detectable APs. Thus, even within the intact node, the superior and inferior regions exhibited visibly different firing patterns.

**Figure 2.**
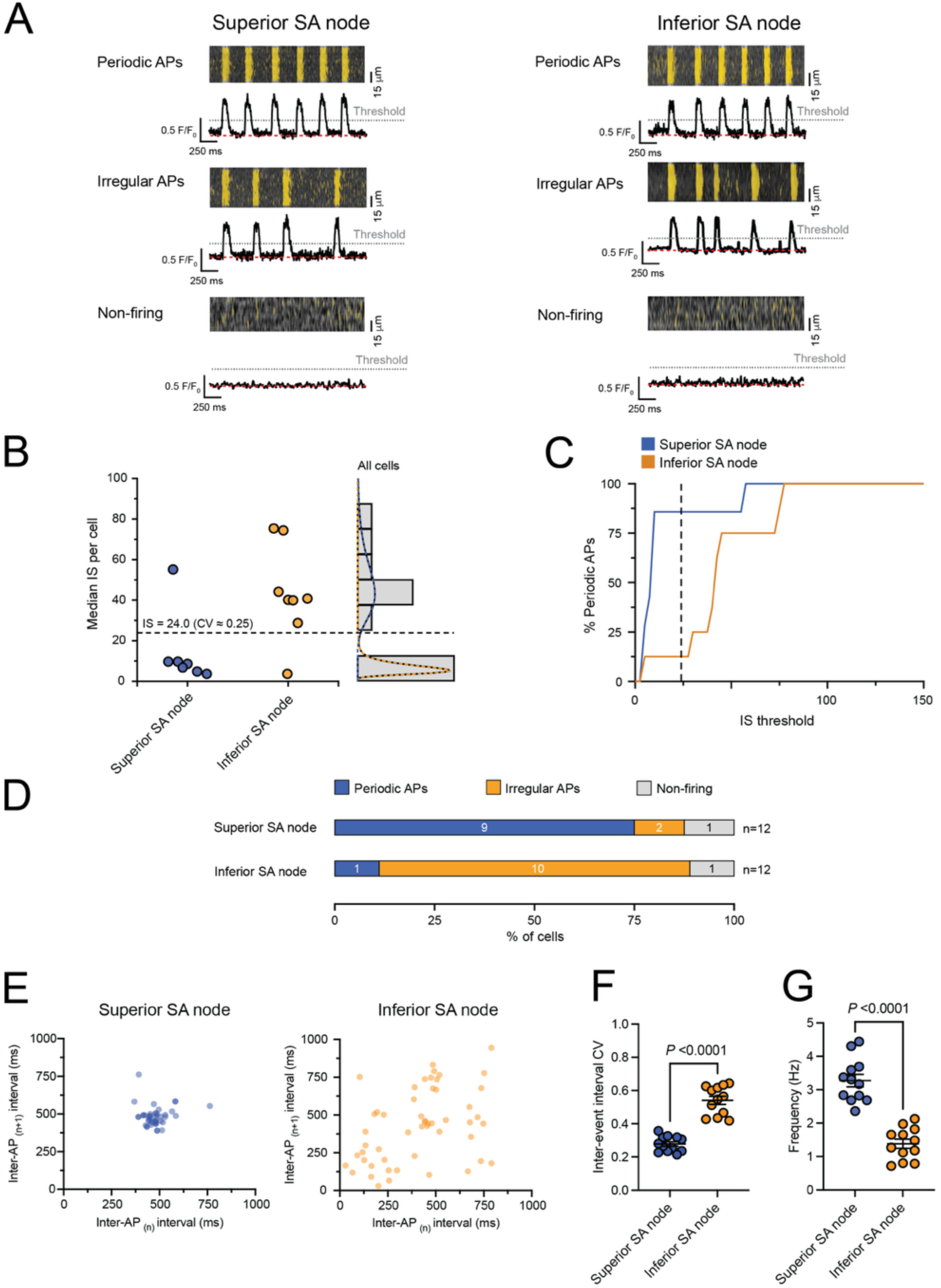
Firing mode shifts from periodic to irregular along the superior–inferior axis. (A) Representative periodic, irregular, and non-firing traces with detected-event images for each pole. (B) Median irregularity score (IS) per recording by pole. Dashed line, classification threshold IS = 24.0 (CV = 0.25). (C) Sensitivity of the classification: percentage of recordings scored as periodic as a function of the IS threshold for each pole. Dashed line, threshold used. (D) Proportions of recordings (*N =* 12) classified as periodic, irregular, and non-firing: superior, 9/2/1; inferior, 1/10/1. (E) Inter-AP interval Poincaré plots (interval n+1 against interval n) for each pole. (F) Inter-event interval CV; higher in the inferior node (*P* < 0.0001). (G) Firing frequency: superior = 3.27 ± 0.19 Hz; inferior = 1.38 ± 0.14 (*P* < 0.0001). Bars show means ± SEM. *P*-values are shown above each comparison. Each circle represents one animal; *N* denotes the number of mice.

Quantifying beat-to-beat regularity requires an explicit, reproducible criterion rather than a qualitative judgment. We therefore scored beat-to-beat regularity with an irregularity score (IS) (Telgkamp *et al*., 2002; Zanella *et al*., 2014). For each pair of successive intervals, we took the absolute change in interval as a percentage of the preceding interval, summarized per cell by its median. IS has two advantages over the coefficient of variation (CV) for this purpose: it is a *local* measure, sensitive to beat-to-beat jitter rather than to slow drift in rate across a recording, and it is not inflated by a cell that is regular in short runs but exhibits a change in mean rate over time.

The distribution of median IS across all recorded cells was bimodal in log space and well described by a two-component Gaussian mixture (**Figure 2B**). We therefore did not impose an arbitrary cutoff, instead placing the boundary between the two populations at the crossing point of the fitted components, IS = 24.0, equivalent to an interval CV of ∼0.25. Cells below this boundary were classified as periodic, cells at or above it as irregular, and cells with fewer than three detected APs as non-firing. The proportion of periodic cells in each pole changed little for thresholds between roughly 10 and 30 (**Figure 2C**), indicating that the classification is stable across a wide range of thresholds and the regional difference does not depend on a precise boundary.

The gradient was clear and separated the poles almost completely. In the superior node, 75% of recordings (9/12) were periodic; of the remaining recordings, two were irregular, and one was non-firing. The pattern was inverted in the inferior node, where 83.3% of recordings (10/12) were irregular and only 8.3% (1/12) were periodic; the remaining recording was non-firing (*P* = 0.0028, periodic vs. non-periodic; Fisher exact test) (**Figure 2D**). Median IS per recording was correspondingly low and tightly grouped in the superior node and high and dispersed in the inferior node (**Figure 2B**). The interval structure underlying this classification was visible directly in the interval Poincaré plots (**Figure 2E**), where superior cells formed a compact cloud near the diagonal in which each interval closely predicted the next. Inferior cells were scattered broadly across the plane and included both very short and very long successive intervals. Consistent with this classification, the inter-event interval CV was higher in the inferior node (*P* < 0.0001) (**Figure 2F**), and firing frequency was lower (1.38 ± 0.14 vs. 3.27 ± 0.19 Hz; *P* < 0.0001) in the inferior node (**Figure 2G**).

These in-tissue measurements agree with previously reported values for isolated SA node myocytes. Grainger *et al*. (2021) found the same three features in dissociated cells sorted by their region of origin: a higher proportion of periodic firing among superior myocytes, faster firing rates in superior compared with inferior cells, and AP waveforms that did not differ between the two populations. That the intact node reproduces all three is a useful cross-validation in both directions. It argues that the regional phenotype is intrinsic to the myocytes rather than imposed solely by the tissue context in which they are recorded, and it indicates that our line-scan classification is measuring the same underlying biology as current-clamp recordings from dissociated cells, despite the very different preparation and readout.

By an explicit, data-derived criterion, then, the superior node fires periodically and the inferior node does not, while the APs themselves are indistinguishable between poles. The regional difference lies in the *timing* of beats rather than in the machinery that generates them, placing the origin of that difference upstream of the AP, within the processes that determine when threshold is reached.

### Subthreshold noise is larger, more spatially fragmented, and more complex in the inferior node

If SVFs are just thermal jitter, they should be small, uniform, and featureless. If instead they are structured biological signals generated by the cell’s own machinery, they should carry statistical fingerprints and could plausibly participate in setting the rhythm. We therefore characterized the diastolic subthreshold signal—the continuous trace between beats, with detected APs removed (**Figure 3A**)—along four axes: amplitude, timescale, spatial extent, and complexity.

**Figure 3.**
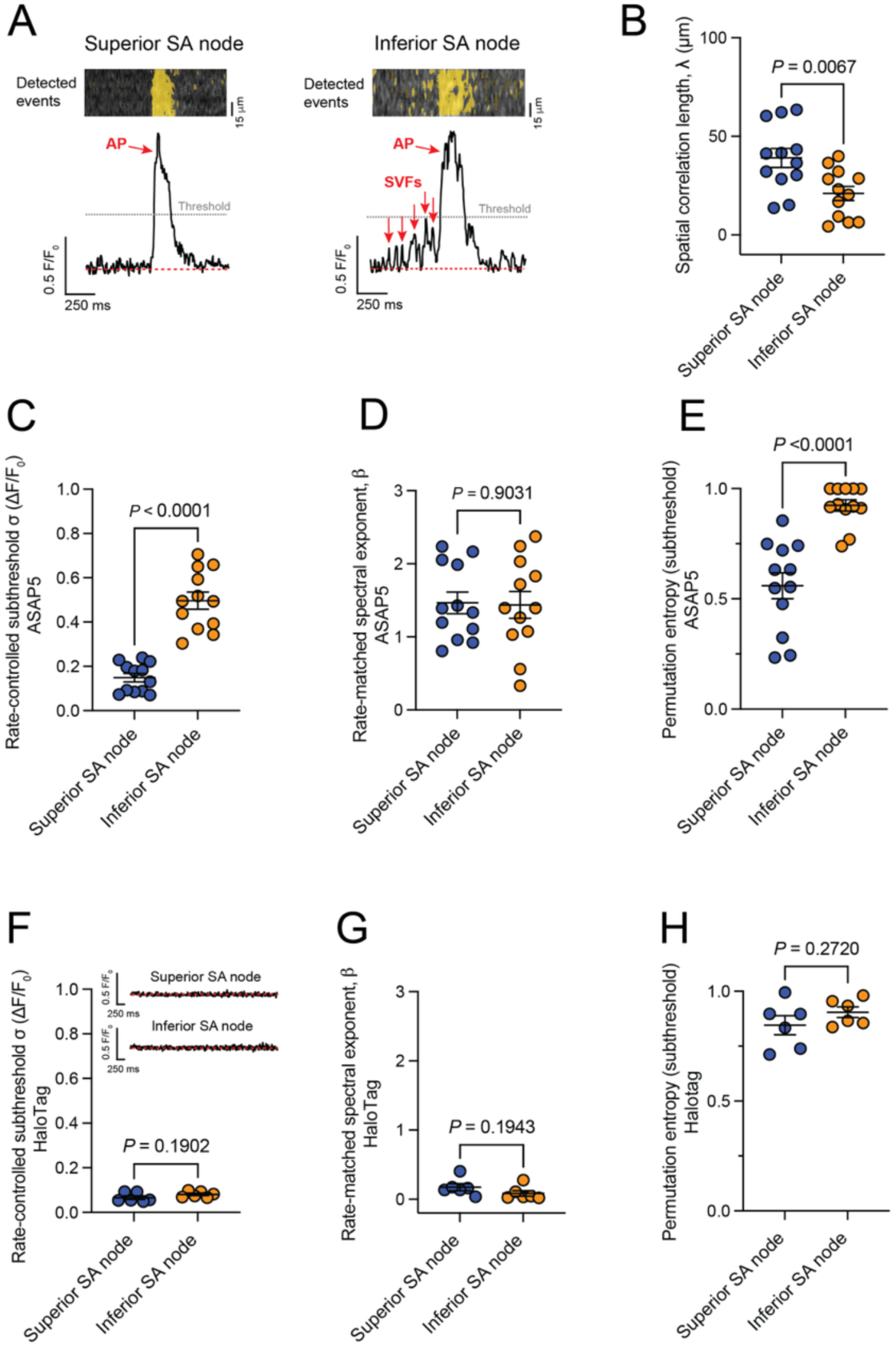
Subthreshold noise is larger, spatially more fragmented, and more complex in the inferior node, whereas the superior–inferior gradient is absent in HaloTag controls. (A) Representative diastolic subthreshold segments (masked line-scan image and ΔF/F_0_ trace) for each pole, with the AP-detection threshold and individual subthreshold voltage fluctuations (SVFs) marked. (B) Spatial correlation length λ (µm) of the SVFs. Superior = 39.05 ± 4.83 µm; inferior = 21.05 ± 3.58 µm (*P* = 0.0067). (C) Rate-controlled subthreshold σ (ΔF/F₀), measured in a fixed-length, ramp-subtracted diastolic window (see Methods). Superior = 0.15 ± 0.02 ΔF/F₀; inferior = 0.49 ± 0.04 ΔF/F₀ (*P* < 0.0001; *N* = 12 for both). (D) Rate-matched aperiodic spectral exponent β of the diastolic PSD (P(f) ∝ f^−β^), with the beat fundamental and harmonics removed. Superior, β = 1.47 ± 0.15; inferior, β = 1.44 ± 0.18 (*P* = 0.9031), placing the subthreshold signal between pink (β = 1) and Brownian (β = 2) noise and excluding a shot-noise–dominated measurement (β ≈ 0). (E) Permutation entropy of the subthreshold ASAP5 signal. Superior = 0.56 ± 0.06; inferior = 0.92 ± 0.03 (*P* < 0.0001). (F) Rate-controlled subthreshold σ (ΔF/F₀) measured from the HaloTag-JFX554 reference signal in superior and inferior SA node regions. Insets show representative HaloTag traces from each region (*P* = 0.1902). (G) Rate-matched spectral exponent β of the HaloTag-JFX554 signal (*P* = 0.1943). (H) Permutation entropy of the HaloTag-JFX554 signal (*P* = 0.2720). Bars show means ± SEM. *P*-values are shown above each comparison. Each circle represents one animal; *N* denotes the number of mice.

We quantified fluctuation amplitude as σ, the standard deviation of the diastolic trace about its own local trend. Because both the diastolic depolarization ramp and the duration of diastole vary with firing rate, and both inflate a naive standard deviation, σ was measured in a fixed-length window placed at a fixed phase of diastole with the linear ramp removed by least squares (see Materials and Methods).

Two measures of subthreshold activity are available in these recordings, and they are built on opposite principles. σ is a continuous statistic of the whole diastolic trace, computed without reference to any detection threshold or event model, whereas the frequency of discrete subthreshold events (**Figure 1D**) requires detection, amplitude classification, and a duration criterion, and uses none of the windowing, ramp subtraction, or rate control that σ depends on. Both report more subthreshold activity in the inferior node. A regional conclusion supported by measures with so little shared machinery is not readily attributable to an artifact of either pipeline.

The two measures are not redundant, and the way in which they differ is itself a regional property. Partitioning σ into the variance attributable to detected events and the variance remaining after those events are accounted for shows that detected events contribute 11.8% ± 3.1% of the measured variance in the superior node but 86.9% ± 4.1% in the inferior node (n = 12 recordings per pole). The subthreshold signal therefore has two separable components: the discrete events the classifier resolves, and a slower, correlated background visible only to a continuous measure (σ). Both components are larger in the inferior node, but they are not graded equally: the event-driven component rises far more steeply between poles than the continuous component. Superior subthreshold activity is predominantly continuous background; inferior subthreshold activity is predominantly discrete. The two measures are thus close to independent in the superior node and substantially overlapping in the inferior node, which is why they agree on direction without being interchangeable.

We take σ as the primary noise variable for three reasons: 1) It is not conditioned on a detection criterion; any event-based measure discards fluctuations that fall below its threshold, so a region whose events are individually smaller loses proportionally more of them and can appear quieter than it is, whereas variance counts all of them. 2) It is model-free, requiring no assumptions about event shape, duration, or amplitude distribution. 3) It aligns directly with the underlying theory; the threshold model is built around overall fluctuation amplitude—the total disturbance acting on the system regardless of its underlying components. In contrast, an event count incorrectly assumes the noise consists of discrete, individually resolvable units. Event frequency is reported alongside it as an independent confirmation of the regional gradient.

Measured in this way, σ was markedly larger in the inferior node: 0.49 ± 0.04 ΔF/F_0_ vs. 0.15 ± 0.02 ΔF/F_0_ in the superior node (*P* < 0.0001) (**Figure 3C**). The same regional difference was present regardless of how the diastolic trace was processed (**Table 1**). In a paired subset of preparations analyzed under both conventions, σ was higher in the inferior node whether measured across the whole diastolic interval with no correction (2.31 ± 0.77-fold, *P* = 0.0011) or in the fixed-length, ramp-subtracted window used throughout (2.64 ± 1.04-fold, *P* = 0.0016). Because the uncorrected measure retains both the depolarization ramp and the full length of diastole, the persistence of the gradient there excludes either as its source. And because rate control widens rather than narrows the difference, regional differences in firing rate and diastolic duration, if anything, mask the difference rather than produce it. SVF amplitude was itself comparable between poles. Both quantities are drawn from the same trace and the same F_0_, so an error in F_0_ would scale them together, and a difference confined to σ cannot arise from the baseline. The subthreshold signal is thus regionally graded rather than uniform, running in parallel with the known energetic gradient.

**Table 1.** Regional comparison of subthreshold voltage noise across processing methods. Subthreshold noise amplitude, σ, was quantified in superior and inferior SA node recordings using two processing variants: (1) the uncorrected whole diastolic interval and (2) a fixed 100-ms diastolic window after removal of the local linear ramp. The rate-controlled measurement (2) was designated the primary analysis and is the σ used throughout. Values are ΔF/F_0_, expressed as means ± SD (one value per recording; N = 5 paired recordings per pole). These are the subset of preparations in which both processing conventions could be applied to the same recordings and are drawn from the larger baseline cohort shown in **Figure 3C**. Group means therefore differ somewhat from that figure, and the comparison here is paired within preparation rather than pooled. Inf/Sup ratios are the means ± SD of the individual inferior-to-superior ratios rather than the ratio of the group means. Regional *P*-values are from two-tailed paired t-tests. A third variant, whole-diastole σ after 500 ms rolling-median detrending, was not included. The trend is estimated from all frames, including APs, whereas σ is computed on diastolic frames only, so the subtracted trend is biased upward near each AP and inflates the diastolic residual in proportion to firing rate. The regional difference is present in the fully uncorrected measurement and widens under rate control; therefore, it cannot be attributed to the diastolic depolarization ramp or to regional differences in firing rate or diastolic duration.

| Processing variant<br>(analyzer field) | Superior $\sigma$ | Inferior $\sigma$ | Ratio Inf/Sup | P<br>(regional) |
| --- | --- | --- | --- | --- |
| Whole diastole, uncorrected<br>(mean $\sigma_{\text{raw}}$) | $0.21 \pm 0.08$ | $0.46 \pm 0.09$ | $2.31 \pm 0.77$ | $p=0.0011$ |
| Fixed 100 ms window, per<br>window linear ramp removed<br>(mean $\sigma_{\text{rate-controlled}}$) | $0.16 \pm 0.06$ | $0.39 \pm 0.06$ | $2.64 \pm 1.04$ | $p=0.0016$ |

We next asked about the timescales of fluctuations using the power spectral density, a decomposition of the signal into the power it carries at each frequency. Featureless ‘white’ noise, like radio static, has flat power across frequencies. Biologically structured ‘colored’ noise has more power at slow timescales, following a power law P(f) ∝ f^−β^. A larger exponent β means more slow, correlated structure and more memory. Because firing rate itself distorts the spectrum, β was computed over a rate-matched band with the beat frequency and its harmonics removed (see Materials and Methods). Both poles gave β ≈ 1.5, with no detectable regional difference (superior 1.47 ± 0.15, inferior 1.44 ± 0.18; *P* = 0.9031) (**Figure 3D**). This places the subthreshold signal between pink noise (β = 1) and Brownian noise (β = 2). Photon shot noise is white by definition and yields β ≈ 0, and because a white component dilutes the fitted exponent in proportion to the variance it contributes, an exponent of 1.5 is incompatible with a shot-noise–dominated measurement. Thus, shot noise cannot account for more than a minor fraction of the variance we report. The two poles therefore share the same noise color but differ substantially in noise amplitude.

The fact that the two poles share an exponent is informative beyond ruling out shot noise, because it also constrains how much of the regional difference could be passive. The membrane behaves as a low-pass filter whose corner frequency, 1/(2πR_in_C_m_), is set by input resistance, so a cell with higher input resistance filters more of the high-frequency content and also converts a given ionic current into a larger voltage deflection. Were the σ gradient principally a matter of higher inferior input resistance, the corner would sit at a lower frequency in that pole, and a fixed-band exponent would differ between poles. It does not, suggesting that the regional difference reflects the subthreshold signal itself and not merely how strongly each membrane expresses it.

We then measured the spatial correlation length λ, how far apart two points along the scan line can be while still fluctuating together. A long λ means large patches move in concert; a short λ means that the tissue breaks into small, independent domains. λ was longer in the superior node (39.05 ± 4.83 µm) than in the inferior node (21.05 ± 3.58 µm; *P* = 0.0067) (**Figure 3B**). Superior subthreshold activity is therefore spatially coherent over longer distances than inferior activity, which is more fragmented. Because SA node myocytes are long (on the order of 100 µm) and thin (on the order of 10 µm) (Reddy *et al*., 2022), and their orientation relative to the scan line is not controlled, the absolute value of λ reflects a mixture of within-cell and between-cell correlation and should not be read as a purely intercellular length. Moreover, because the same sampling approach was applied to both poles, the regional difference is unlikely to arise from cell geometry alone unless myocyte orientation relative to the scan line differs systematically between regions. Photon shot noise between neighboring positions is uncorrelated, meaning its correlation length is limited to the scale of a single pixel; thus, this measure provides a second, independent argument that the signal is biological.

Finally, we measured the permutation entropy of the subthreshold trace. Entropy here is a model-free score of how unpredictable a signal’s moment-to-moment sequence is. The method looks at short runs of consecutive samples, asks which up-down ordering pattern each run takes, and counts how evenly the possible patterns are used. A smooth, predictable signal reuses a few patterns and scores low, near 0; a complex, disordered signal spreads across all patterns and scores high, near 1. Superior traces scored 0.56 ± 0.06 and inferior traces 0.92 ± 0.03 (*P* < 0.0001) (**Figure 3E**). Thus, the inferior subthreshold signal is not only larger but genuinely less predictable in its ordinal structure.

On all four measures, the fluctuations behave like endogenous, structured signals rather than thermal or photonic jitter: colored, spatially organized, regionally graded, and complex. All four measures also converge on one account of what separates the poles. The subthreshold signal has a slow, correlated component that σ and β report and a faster, discrete component classified in **Figure 1D**. Moreover, the two are graded differently: superior activity is predominantly the continuous background; inferior activity is predominantly the discrete events. Event frequency, variance partition, and permutation entropy each rest on different assumptions and each return that same picture. Both components are larger in the inferior node, but the discrete component rises far more steeply.

To test whether the superior–inferior differences in subthreshold ASAP5 dynamics arise from membrane voltage rather than from the optics, the tissue, or residual contractile motion, we recorded a voltage-insensitive reference. HaloTag is a self-labeling protein that forms a covalent bond with a synthetic fluorophore, here the photostable dye JFX554, yielding a single fluorescent molecule whose brightness does not report membrane potential. We used the same reporter for this purpose in the intact node and ventricular myocytes previously (Muñoz *et al*., 2026; Rhana *et al*., 2026). It registers only the nonspecific fluorescence fluctuations that every optical recording in this preparation carries. HaloTag-JFX554 fluctuations, analyzed identically, were smaller than those of ASAP5 and regionally uniform (σ: superior 0.06 ± 0.01, inferior 0.08 ± 0.01; *P* = 0.1902; **Figure 3F**). ASAP5 was 1.2-fold brighter than HaloTag-JFX554, so the control carried the higher shot-noise floor of the two. The spectral exponent β sat near zero in both poles (superior 0.17 ± 0.05, inferior 0.08 ± 0.04; *P* = 0.1943; **Figure 3G**), the signature of shot noise rather than of a signal with temporal structure. Permutation entropy was correspondingly high and likewise indistinguishable between regions (superior 0.84 ± 0.04, inferior 0.90 ± 0.02; *P* = 0.2720; **Figure 3H**). Neither the amplitude gradient nor the temporal-complexity gradient seen with ASAP5 survives in a voltage-insensitive fluorophore imaged in the same tissue under the same illumination. The regional differences in ASAP5 signals are therefore properties of membrane voltage. They also settle the assumption carried by the opening paragraph: with contraction arrested by blebbistatin, residual motion adds no structured fluctuation to the fluorescence record.

### Diastolic coherence peaks at an intermediate level of noise: stochastic resonance

We now have two regional gradients that run in parallel: periodic-versus-irregular firing (**Figure 2**) and low-versus-high subthreshold noise (**Figure 3**). Our central hypothesis is that these are causally linked through stochastic resonance: a counterintuitive phenomenon, well known in physics and neuroscience, in which a *moderate* amount of noise *improves* a system’s ability to produce a rhythmic output (Bezrukov & Vodyanoy, 1995; Pérez-Cervera *et al*., 2023). With too little noise, a weak periodic drive never crosses the firing threshold, and the cell stays quiet; with too much noise, the timing is swamped and the rhythm scatters. Only at an intermediate “Goldilocks” level does noise reliably nudge the drive across threshold on the beat, sharpening the rhythm. A testable prediction of this model is that rhythm quality plotted against noise level should trace an inverted-U, peaking at an intermediate optimum.

To test this hypothesis, we first need a measure of how well neighboring cells beat in step. For this, we used the Kuramoto order parameter R, a standard index of synchrony (Breakspear *et al*., 2010). Each cell can be represented as a point moving around a circle, completing one revolution per heartbeat cycle, so that its position on the circle gives its phase within the cycle. R measures how tightly those points are clustered at a given instant. A value near 1 indicates that the cells occupy nearly the same phase, and a value near 0 that their phases are dispersed around the circle. We computed R over the diastolic interval, defined here from the end of one AP through the upstroke of the next, so that the measure spans both the subthreshold approach to threshold and the moment at which each cell is activated (**Figure 4A**). Coherence is therefore assessed across the full interval over which cells commit to the next beat.

**Figure 4.**
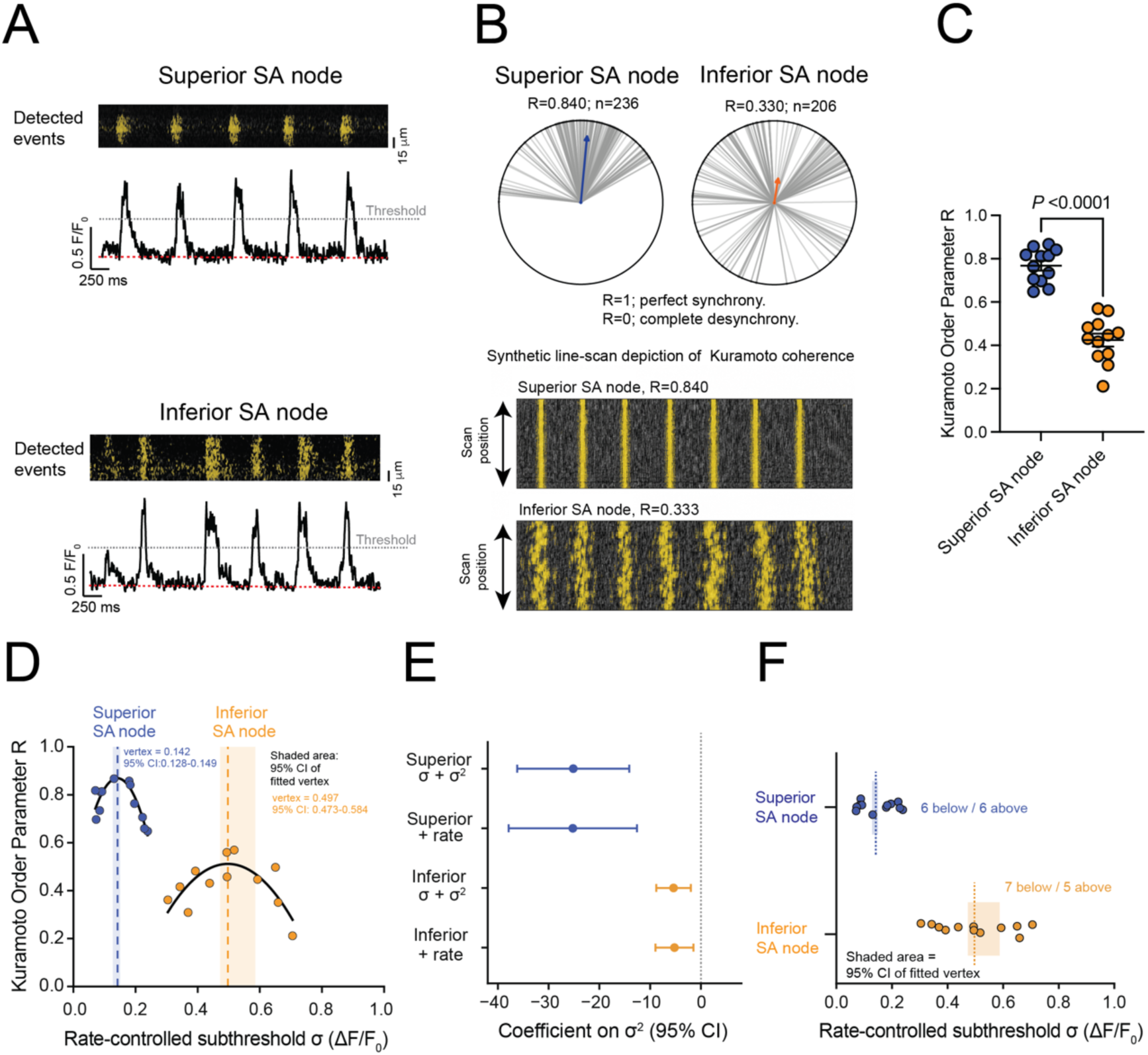
Diastolic coherence peaks at an intermediate level of subthreshold noise (stochastic resonance). (A) Representative baseline line-scan images and ΔF/F₀ traces from the superior and inferior SA node. (B) Phase-vector plots showing diastolic phase across valid spatial columns; resultant length = Kuramoto R. Superior: R = 0.84 (n = 236 columns); inferior: R = 0.33 (n = 206 columns). (C) Group diastolic Kuramoto R: superior, 0.78 ± 0.02; inferior, 0.42 ± 0.03 (*P* < 0.0001; *N* = 12 per pole). (D) Kuramoto R versus rate-controlled subthreshold σ, with quadratic fits within each pole. Fitted vertices: superior, σ = 0.142 (95% CI 0.128–0.149); inferior, σ = 0.497 (95% CI 0.473–0.584). Shaded bands denote vertex 95% CIs. (E) Curvature remained significantly negative without and with firing-rate adjustment: superior, −25.15 (95% CI −36.19 to −14.10; *F*(1,9) = 26.53, *P* = 0.0006) and −25.24 (95% CI −37.89 to −12.59; *F*(1,8) = 21.17, *P* = 0.0018); inferior, −5.39 (95% CI −8.80 to −1.98; *F*(1,9) = 12.78, *P* = 0.0060) and −5.21 (95% CI −8.96 to −1.46; *F*(1,8) = 10.26, *P* = 0.0126). Adding firing rate did not alter curvature (superior Δ = −0.09, *P* = 0.962; inferior Δ = 0.18, *P* = 0.659). (F) Both limbs of each fitted curve were populated: 6 below/6 above the vertex in the superior node and 7 below/5 above in the inferior node. Blue, superior; orange, inferior. Bars show means ± SEM. Each circle represents one animal; *N* denotes the number of mice.

**Figure 4B** depicts this graphically for one representative recording per pole, drawing each cell’s diastolic phase as an arrow on a circle, with the length of their average (the resultant) equal to R. In the representative superior recording, the arrows bunch tightly (R = 0.84, n = 236 columns); in the representative inferior recording, they fan out around the circle (R = 0.33, n = 206). A paired synthetic line-scan panel translates these two R values back into the visibly tighter versus more scattered beat patterns they produce.

On average, diastolic coherence was high in the superior SA node (R= 0.78 ± 0.02) and low in the inferior node (R= 0.42 ± 0.03; *P* < 0.0001) (**Figure 4C**). The superior node therefore behaves as a tightly phase-locked oscillator and the inferior node as a loosely coupled, noise-dominated population, matching the firing-mode gradient of **Figure 2**.

The decisive test is the relationship between coherence and noise level. Plotting diastolic R against subthreshold noise amplitude σ and fitting a quadratic within each pole, we found that coherence followed the predicted inverted-U relationship in both: R was maximal at an intermediate σ and fell when noise was either lower or higher (**Figure 4D**). The curvature was significant in each pole taken separately, with a negative coefficient on σ^2^ (superior: −25.15; inferior: −5.39) and a 95% confidence interval (CI) that excluded zero (superior: 95% CI −36.19 to −14.10; inferior: 95% CI −8.80 to −1.98) (**Figure 4E**). Because σ and firing rate covary, we refitted the model with firing rate as a covariate. The curvature was essentially unchanged in both the superior (coefficient, −25.24; 95% CI −37.89 to −12.59) and inferior (coefficient, −5.21; 95% CI −8.96 to −1.46) nodes, and the change in curvature attributable to rate was negligible (superior Δ = −0.09, *P* = 0.962; inferior Δ = 0.18, *P* = 0.659). The non-monotonic dependence of coherence on σ is therefore not a firing-rate effect in disguise.

The fitted vertices for the two poles were also well constrained within the experimentally sampled σ ranges, with σ = 0.142 (95% CI 0.128–0.149) in the superior node and σ = 0.497 (95% CI 0.473–0.584) in the inferior node (**Figure 4D**). Recordings populated both sides of each vertex, with six below and six above in the superior node and seven below and five above in the inferior node (**Figure 4F**). Thus, the vertices are supported by observations on both the ascending and descending limbs rather than arising from extrapolation beyond the sampled data. The mirror-image test agreed: CV-IBI (i.e., beat-to-beat interval variability), for which lower values indicate a more regular rhythm, traced a U against σ and was minimized near the same intermediate level in each pole (**Supplementary Figure 1**); it was also lower overall in the superior node than in the inferior node. Coherence and interval variability therefore respond to σ as mirror images, as resonance requires.

Because the Kuramoto order parameter is bounded between 0 and 1, we additionally tested whether the nonlinear relationship was robust to modeling R on a logit-transformed scale. Significant negative curvature was retained in both the superior (σ^2^ coefficient = −145.37, 95% CI −210.78 to −79.95, *F*(1,9) = 25.27, *P* = 0.0007) and inferior (σ^2^ coefficient = −23.44, 95% CI −38.22 to −8.67, *F*(1,9) = 12.88, *P* = 0.0059) nodes. The fitted vertices were nearly unchanged from the primary analysis (superior, σ = 0.143; inferior, σ = 0.495), and including firing rate did not significantly alter curvature in either region (*P* = 0.958 and *P* = 0.665 for the change in curvature in superior and inferior nodes, respectively). Thus, the inverted-U relationship was robust to the bounded nature of R, was not explained by firing rate, and was supported within the experimentally sampled range (**Table 2**).

**Table 2.**
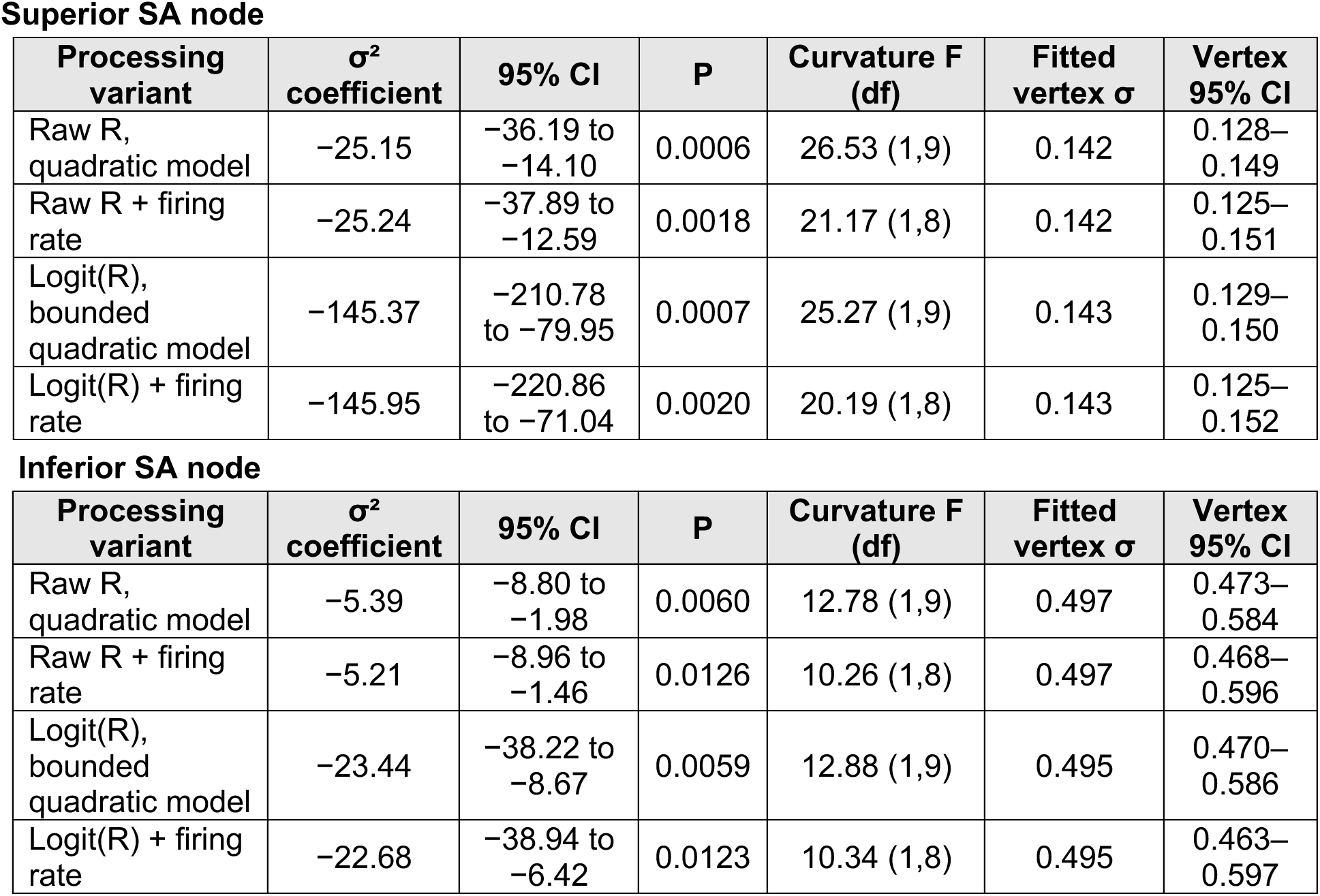
Sensitivity analysis of the nonlinear relationship between subthreshold noise and Kuramoto coherence. Values are quadratic σ^2^ coefficients with 95% CIs from models fitted separately in the superior and inferior SA node (N = 12 recordings per region). Raw-R models are the primary analysis shown in **Figure 4**. Because the Kuramoto order parameter R is bounded between 0 and 1, logit(R) models were used as a bounded-response sensitivity analysis. Models were fitted with and without firing rate as a covariate. Curvature F statistics are partial F tests for the quadratic σ^2^ term; degrees of freedom are shown in parentheses. Fitted vertices were calculated from the quadratic terms, and vertex 95% CIs were obtained by nonparametric bootstrap resampling of recordings. Significant negative curvature persisted in both regions after logit transformation and after adjustment for firing rate. Adding firing rate did not significantly alter curvature: superior, raw R Δσ^2^= −0.09 (95% CI −4.26 to 4.08, *P* = 0.962) and logit(R) Δσ^2^= −0.58 (95% CI −25.29 to 24.13, *P* = 0.958); inferior, raw R Δσ^2^ = +0.18 (95% CI −0.73 to 1.09, *P* = 0.659) and logit(R) Δσ^2^= +0.77 (95% CI −3.16 to 4.70, *P* = 0.665).

To determine whether regional differences in coherence could be explained by systematic propagation delays across the tissue, we recalculated Kuramoto R after correcting for the spatial lag in activation timing between recording columns. Lag correction produced essentially no change in coherence in either region. In the superior SA node, R was 0.7261 ± 0.1240 before correction and 0.7276 ± 0.1242 after correction, with a mean change of only 0.0015. Similarly, inferior R was 0.4331 ± 0.1492 before correction and 0.4334 ± 0.1491 after correction, with a mean change of only 0.0003. Thus, the marked difference in coherence between the superior and inferior regions persisted after removal of systematic spatial activation delays. Because such delays represent the component of the signal most directly associated with electrical propagation through intercellular coupling, including gap junction-mediated conduction, these results argue against differences in propagation or gap junction coupling as the primary explanation for the regional coherence pattern. Instead, the persistence of the difference after lag correction supports the interpretation that local subthreshold voltage dynamics, rather than a conduction delay, are the major determinant of the observed regional differences in coherence (**Table 3**).

**Table 3.** Effect of spatial lag correction on Kuramoto coherence in the superior and inferior SA node. Raw and lag-corrected Kuramoto order parameter (R) values are presented as means ± SD with 95% CIs. Lag correction accounted for systematic spatial differences in activation timing across recording columns before recalculating R. Values were essentially unchanged after correction in both regions, indicating that the regional difference in coherence was not explained by spatial propagation delay.

| Region | Raw R, mean ± SD<br>(95% CI) | Lag corrected R,<br>mean ± SD<br>(95% CI) | Mean ΔR<br>(95% CI) |
| --- | --- | --- | --- |
| Superior<br>SA node<br>(N=9) | 0.7261 ± 0.1240<br>(0.6308 to 0.8214) | 0.7276 ± 0.1242<br>(0.6322 to 0.8231) | +0.0015<br>(0.0005 to 0.0026) |
| Inferior<br>SA node<br>(N=9) | 0.4331 ± 0.1492<br>(0.3184 to 0.5477) | 0.4334 ± 0.1491<br>(0.3188 to 0.5480) | +0.0003<br>(-0.0005 to 0.0011) |

Two features of these curves matter for what follows. *First*, the poles do not share a single relationship. Each traces its own inverted-U curve, with the superior vertex near σ ≈ 0.15 and the inferior near σ ≈ 0.5, and with the superior curve reaching a substantially higher peak coherence. Their baseline σ values (**Figure 3C**) place each pole close to its own vertex rather than at different points along one common curve. What separates them is therefore not distance from a shared optimum but the height of the optimum each can reach, consistent with the poles differing in the strength of their periodic drive as well as noise amplitude, a distinction addressed directly in **Figures 6 and 7**. *Second*, because the poles differ in drive, coupling and diastolic input impedance, a pole is properly compared against its own optimum rather than against a common curve; accordingly, we fit and interpret them separately throughout.

There is a subtlety worth making explicit, because it is the literal meaning of ‘order from noise’. The order that resonance creates lives in the rhythm, the regular spacing of beats, indexed by low CV-IBI and high R, not in the underlying fluctuations, which remain complex. The inferior node has both the larger σ and the higher subthreshold permutation entropy (**Figure 3E**), yet its coherence is nonetheless maximal at an intermediate rather than a minimal noise level. A noisier moment-to-moment signal, at the right amplitude for that pole, yields a more ordered beat. To our knowledge, this is the first measurement of the stochastic resonance inverted-U relationship in membrane potential within intact pacemaker tissue, and the first in which the fluctuations are endogenous rather than injected, with each pole occupying its own metabolically established optimum.

### Spatial organization of the pacemaker resonance

There is a spatial counterpart to the regional difference in coherence. We asked what site within the field leads the beat, and whether that leading site holds its position, reasoning that a more coherent, longer-range–correlated pole should also hold a steadier leading pacemaker.

It did. Leading-site stability—the fraction of time the same location initiated the beat—was higher in the superior node (17.58% ± 1.77 %) than in the inferior node (9.88% ± 0.68 %) (**Figure 5A**). The superior lead was consistent; the inferior lead wandered. The mean leading-site position was displaced between poles (**Figure 5B**), consistent with the superior region acting as the dominant initiation zone. This steadier lead paralleled the longer subthreshold spatial correlation length of the superior node (39.05 ± 4.83 µm) compared with the inferior node (21.05 ± 3.58 µm) (**Figure 3B**); the pole whose subthreshold signal is spatially coherent over longer distances also anchors its leading site. Coherence is therefore a genuinely tissue-level property. The same metabolic and noise gradient that sets phase coherence also sets where the rhythm is led and how stably, which is what an emergent, spatially organized phenomenon should look like and what cannot be produced as a property of isolated cells.

**Figure 5.**
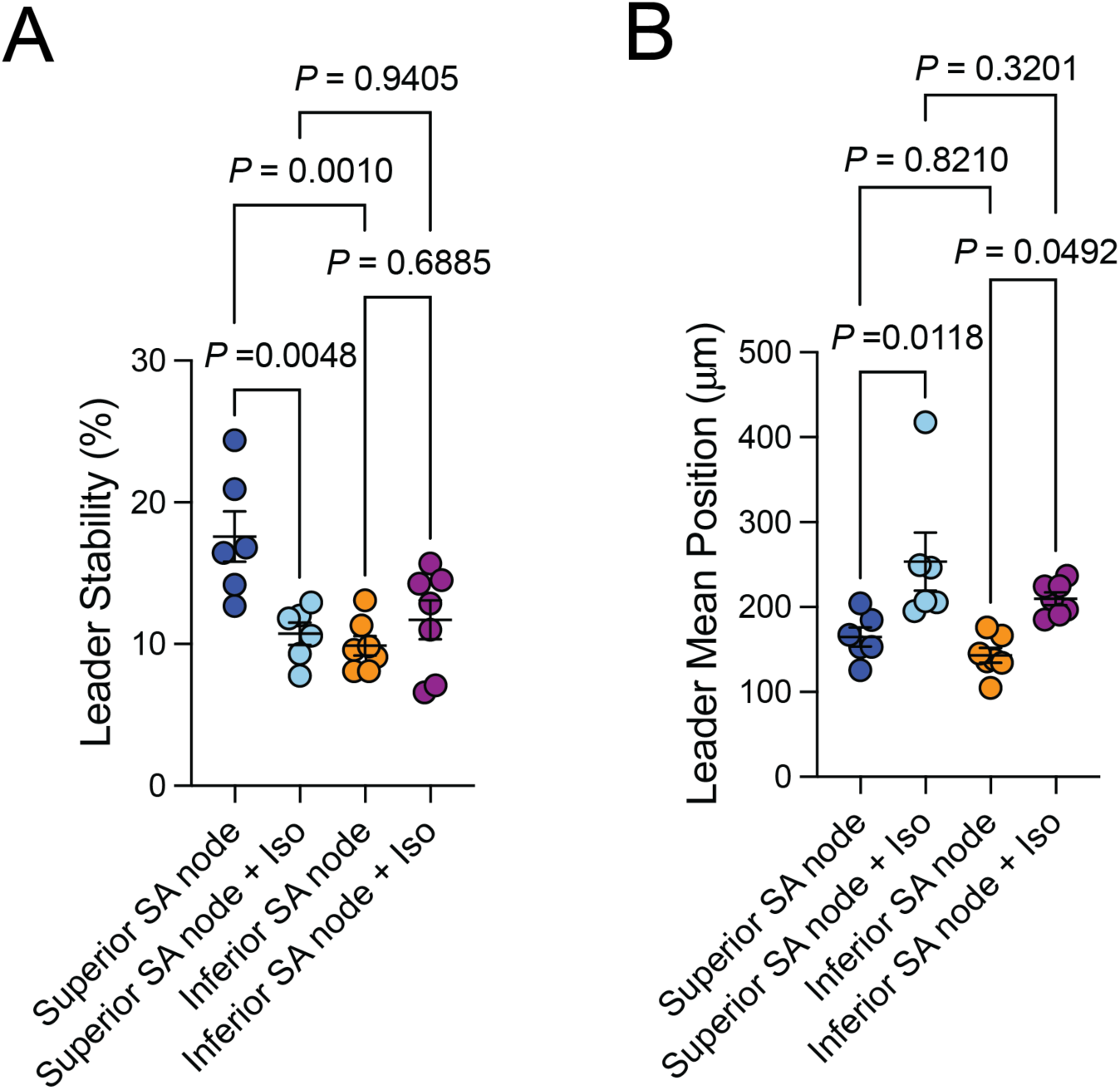
The leading pacemaker site is more stable in the superior node and shifts under β-adrenergic drive. (A) Leader stability (%) across the four groups (superior, superior + isoproterenol, inferior, inferior + isoproterenol) (superior, *N* = 6; inferior, *N* = 7). (B) Leader mean position (µm) across the same four groups. Spatial correlation length λ is shown with the baseline noise measures in Figure 3B. Each point represents one recording. Bars show means ± SEM. Exact *P*-values are shown. Isoproterenol reduces leading-site stability in the superior node and displaces the mean leading position in both poles, reorganizing the spatial structure of the rhythm.

The inverted-U of **Figure 4** makes a testable prediction: if coherence really is a resonance, then moving a preparation along the noise axis should change its rhythm in a direction that depends on where that preparation starts. A pole below its optimum should improve; a pole at or beyond its optimum should not. Two poles beginning at different positions on their own curves therefore provide a built-in test with an unusually specific expected outcome, and one that can be run within single preparations.

Two separable variables set a pole’s position on the resonance surface: the amplitude of the subthreshold noise (σ) and the strength of the periodic pacemaker drive carried by the membrane and Ca^2+^ clocks. Coherence is a surface over both—a resonance ridge in the noise × drive plane—so a manipulation raises or lowers coherence according to which coordinate it moves and in which direction, and a change in drive moves the optimum itself rather than sliding a pole along a fixed curve. Isoproterenol, ivabradine, and PIP_2_ each act predominantly on a different coordinate, and the action of each is interpreted based on that coordinate.

### Isoproterenol moves each pole along its own resonance curve, with opposite consequences

We began with isoproterenol, a β-adrenergic agonist that increases I_f_ and Ca^2+^ cycling and should therefore raise subthreshold noise, pushing each pole rightward along the σ axis. The effect was immediately visible in the raw recordings. **Figure 6A** shows line scans and their extracted traces from the superior and inferior node before and during isoproterenol (100 nM) administration. Beats became faster in both poles, as expected of β-adrenergic stimulation, and the diastolic intervals between them became visibly more ‘agitated’, with subthreshold events both more frequent and larger. In the superior node, which was quiet between beats at baseline, the change was striking: diastole filled with fluctuations that approached the amplitude threshold. The inferior node, already active at baseline, became more so.

**Figure 6.**
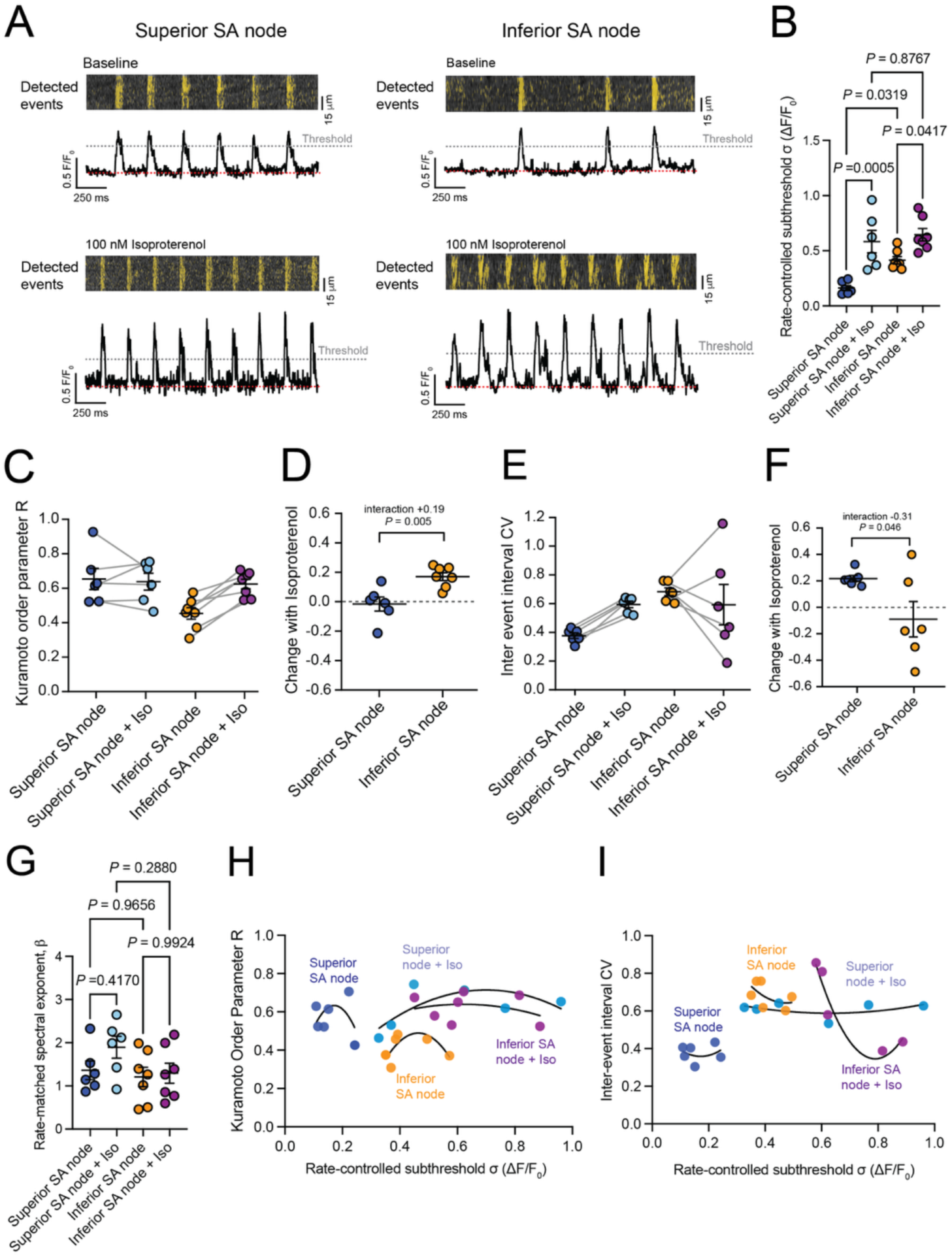
Isoproterenol shifts each pole along its own coherence–noise curve. (A) Representative line-scan images of detected events and corresponding fluorescence traces from the superior and inferior SA node at baseline and after 100 nM isoproterenol. Gray dotted lines indicate the threshold. Red dotted lines indicate baseline. (B) Mean rate-controlled subthreshold σ across the four groups (superior, *N* = 6; inferior, *N* = 7). (C) Kuramoto order parameter R, paired within preparation. (D) Change in R with isoproterenol by pole; pole × drug interaction = +0.19 as estimated by a mixed-effects model (*P* = 0.005). (E) Inter-event interval CV, paired within preparation. (F) Change in inter-event interval CV with isoproterenol by pole; interaction = −0.31 (*P* = 0.046). (G) Rate-matched spectral exponent β. (H) Kuramoto order parameter R plotted against mean rate-controlled subthreshold σ, with quadratic fits. (I) Inter-event interval CV plotted against mean rate-controlled subthreshold σ. Colors indicate superior SA node, superior SA node + isoproterenol, inferior SA node, and inferior SA node + isoproterenol. Exact *P*-values are shown. Bars show means ± SEM. Each circle represents one animal; *N* denotes the number of mice.

These drug experiments were paired within preparation, with each field serving as its own control, and were acquired in a cohort separate from the unpaired baseline survey of **Figures 2–4**. Baseline values therefore differ between the two sets of figures, and every drug effect reported below is assessed within each preparation rather than against the pooled baselines. Quantification confirmed that isoproterenol moved the intended variable, raising σ from 0.16 ± 0.02 to 0.58 ± 0.10 in the superior node (*P* = 0.0005) and from 0.41 ± 0.03 to 0.65 ± 0.05 in the inferior node (*P* = 0.0417) (**Figure 6B**). The displacement was much larger superiorly. After isoproterenol, σ no longer differed between the two poles at all (*P* = 0.8767), so the drug abolishes the regional noise gradient.

Coherence responded differently in the two poles, and that difference is the informative result. In the inferior node, which began with lower coherence, R rose from 0.42 ± 0.03 to 0.62 ± 0.03 (*P* = 0.0022) (**Figure 6C**), while interval variability was unchanged (*P* = 0.7219) (**Figure 6E**). In the superior node, which in these preparations began near its own coherence maximum (**Figure 6H**), added noise produced no improvement in R (0.57 ± 0.04 to 0.62 ± 0.04, *P* = 0.7588) and significantly worsened interval variability (inter-event interval CV, 0.37 ± 0.02 to 0.61 ± 0.02, *P* = 0.0068). Thus, the same drug moving the same coordinate in the same direction improved the rhythm of one pole and degraded that of the other. Testing that difference directly rather than by comparing two separate *P*-values, we found that the pole × drug interaction was significant for both readouts: isoproterenol changed R by +0.19 more in the inferior node than in the superior node (*P* = 0.005) (**Figure 6D**) and changed interval variability by −0.31 more in the inferior node than in the superior node (*P* = 0.046) (**Figure 6F**). The opposing regional response is therefore a formal interaction result, not an inference from two independent comparisons. Lastly, β-adrenergic stimulation did not alter the rate-matched spectral exponent β in either the superior or inferior SA node, indicating that noise color remained unchanged (**Figure 6G**).

This is what a resonance predicts and what a monotonic noise effect does not: a pole below its optimum gains from added fluctuation, and a pole already at its optimum is pushed onto the descending limb of the inverted-U curve, where further noise scatters the timing. Plotting each recording’s coherence against its own σ makes the geometry explicit (**Figure 6H**). Points in the superior node span the vertex and continue well beyond it, tracing the full arc from ascending limb through peak to descent, while points in the inferior node remain concentrated near that pole’s own maximum. The mirror-image relationship holds for interval variability (**Figure 6I**), where the superior trajectory rises away from its minimum as σ increases. The descending limb, which in **Figure 4** could only be inferred from a fit across recordings, is here traversed within preparations.

Three further observations follow. Isoproterenol also flattens the coherence–σ relationship in both poles, so that coherence becomes markedly less dependent on noise amplitude under β-adrenergic drive than at baseline. On the two-coordinate reading, this is expected, since isoproterenol increases drive as well as noise and therefore moves a preparation along the crest of the ridge rather than across it. More broadly, it suggests that noise-dependent coordination is a property of the resting node, one that recedes when catecholaminergic drive supplies coordination directly. And because isoproterenol equalizes σ between the poles while leaving their coherence similar in superior (R = 0.62 ± 0.04) and inferior (R = 0.62 ± 0.02) SA nodes, it simultaneously collapses the regional gradient in both the input and output variable. Third, isoproterenol also reorganizes the spatial structure of the rhythm, reducing leading-site stability in the superior node and displacing the mean leading position in both poles (**Figure 5A, B**). Pushing a pole along its resonance curve therefore moves where and how stably the beat is led, in addition to affecting its average coherence.

This result converges with the autonomic modulation of stochastic resonance reported in isolated cells by Okamura *et al*. (2026), where cholinergic stimulation shifted each cell’s resonance spectrum toward lower frequencies and injected noise then improved firing in the slowed cells. These authors’ basal-state data showed the same dichotomy we report here: injected noise improves rhythmicity in slow-firing cells while degrading it in fast-firing cells. That this dichotomy reappears between anatomically defined poles, in a different species, in intact tissue, and with fluctuations raised endogenously rather than delivered through an electrode argues that the outcome is set by where a cell sits on its own resonance surface rather than by the properties of the applied current.

### Weakening pacemaker drive with ivabradine lowers coherence in the superior node

Isoproterenol probed the noise coordinate of the resonance landscape. The complementary test is to weaken the periodic drive itself—the ‘weak signal’ ingredient of stochastic resonance—and ask whether coherence falls. We used ivabradine, a selective blocker of the funny current (I_f_), the hyperpolarization-activated current carried by HCN4 channels that provides the rhythmic depolarizing push between beats (Bucchi *et al*., 2002; Bucchi *et al*., 2006). Blocking I_f_ should degrade the periodic drive and move cells off their coherence optimum.

Ivabradine (50 µM) narrowed the beat pattern in both poles, as visibly evident in the raw recordings (**Figure 7A**), and the two poles then diverged. Critically, subthreshold σ was unchanged in the superior node (*P* = 0.7136), whereas in the inferior node, ivabradine markedly reduced subthreshold σ from 0.48 ± 0.03 to 0.15 ± 0.01 (*P* = 0.0053) (**Figure 7B**), abolishing the regional noise difference (superior vs. inferior + ivabradine, *P* = 0.2779).

**Figure 7.**
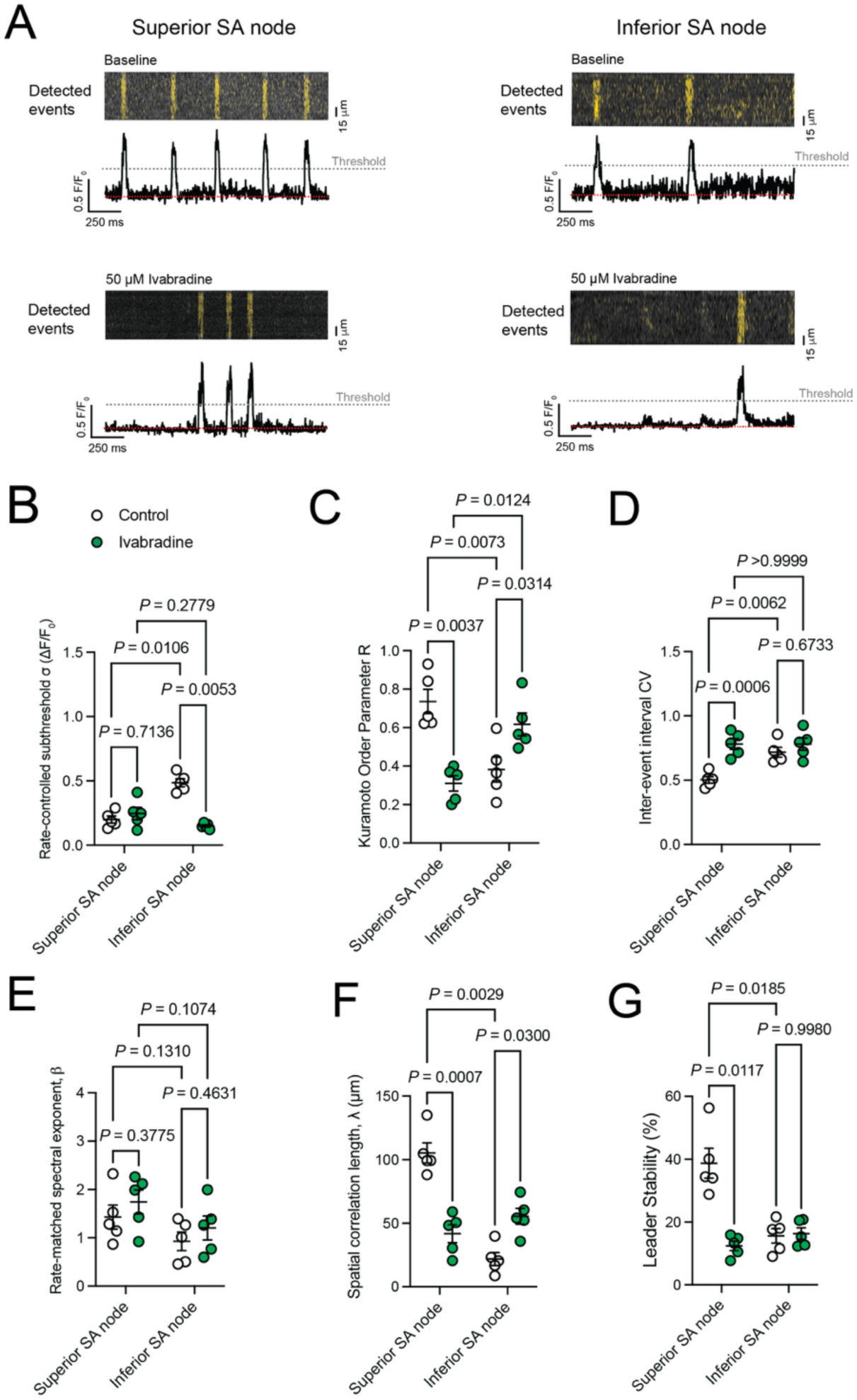
Ivabradine weakens the periodic drive, lowering coherence in the superior node and reducing subthreshold noise in the inferior node. (A) Representative detected-event images and ΔF/F_0_ traces at baseline and during 50 µM ivabradine for each pole. (B) Mean rate-controlled subthreshold σ: unchanged in the superior node (*P* = 0.7136) and markedly reduced in the inferior node (*P* = 0.0053; *N* = 5 for both poles). (C) Kuramoto order parameter R during diastole: reduced superiorly (*P* = 0.0037) and increased inferiorly (*P* = 0.0314). (D) Inter-event interval CV: increased superiorly (*P* = 0.0006) and unchanged inferiorly (*P* = 0.6733). (E) Rate-matched spectral exponent β: unchanged in both poles. (F) Spatial correlation length λ: reduced superiorly (*P* = 0.0007) and increased inferiorly (*P* = 0.0300). (G) Leader stability: reduced superiorly (*P* = 0.0117) and unchanged inferiorly (*P* = 0.9980). Bars show means ± SEM. Exact *P*-values are shown. Each circle represents one animal; *N* denotes the number of mice.

Despite the absence of a change in σ, diastolic coherence in the superior node fell sharply from R = 0.74 ± 0.06 to R = 0.31 ± 0.04 (*P* = 0.0037) (**Figure 7C**). Coherence therefore fell without any movement along the noise axis, distinguishing a loss of drive from a displacement in noise. Movement along the resonance curve in either direction requires σ to change, which it did not. Thus, the superior decoherence is a move down the drive axis. The inferior node behaved differently: rather than falling, diastolic coherence increased (*P* = 0.0314) (**Figure 7C**). A pole whose coherence is already low, and whose drive was weak to begin with, has less to lose when that drive is reduced further.

Interval variability rose in the superior node (*P* = 0.0006), whereas it did not change in the inferior node (*P* = 0.6733) (**Figure 7D**). The spectral character of the fluctuations was preserved throughout. The spectral exponent β was unchanged in both poles (superior, *P* = 0.3775; inferior, *P* = 0.4631) (**Figure 7E**), while the spatial correlation length moved in opposite directions, falling in the superior node (*P* = 0.0007) and rising in the inferior node (*P* = 0.0300) (**Figure 7F**). Weakening the drive therefore rearranged the spatial organization of the subthreshold signal without altering the timescales on which it operates.

Finally, leading site stability collapsed in the superior node from 38.73% ± 4.75% to 12.44% ± 1.49% (*P* = 0.0117), whereas it did not change in the inferior node (*P* = 0.9980) (**Figure 7G**). Notably, ivabradine does not abolish the beat-locked ATP fluctuations we described previously (Muñoz *et al*., 2026); by blocking the voltage current while leaving the metabolic oscillation intact, it isolates the voltage and I_f_ arm of the pacemaker from its energy supply.

Ivabradine therefore dissociated the response between the poles: the superior node lost coherence, rhythm regularity, and leading-site stability without a change in its subthreshold noise, whereas the inferior node lost noise without a corresponding loss of coherence. Weakening I_f_ moved the drive coordinate directly, and in the pole that began with the strongest drive and the highest coherence, it lowered both. Together with the isoproterenol experiments, which manipulated the noise coordinate, **Figures 6–7** provide within-preparation tests of the effects of independently manipulating the two coordinates of the resonance surface.

### PIP_2_ acts differentially across the node, more strongly accelerating firing inferiorly and improving coherence only in the inferior node

The results to this point link a cell’s position on the resonance curve to its subthreshold noise, which we posit is metabolically set (Muñoz *et al*., 2026). This predicts a specific, testable asymmetry: If the inferior node fires poorly because a low ATP ceiling limits an ATP-sensitive subthreshold conductance, then relieving that limitation should improve firing in the energy-poor inferior node far more than in the energy-rich superior node, which is not ATP-limited. The molecular handle is PIP_2_, a membrane lipid that sustains I_f_ (Pian *et al*., 2006; Pian *et al*., 2007). Maintaining PIP_2_ is itself ATP-dependent, so an energy-starved cell loses PIP_2_ support for I_f_, and supplying PIP_2_ directly should bypass that bottleneck. To test this, we applied membrane-permeant diC8-PIP_2_ during two-photon ASAP imaging and compared the two poles (**Figure 8**). As shown in **Figure 8A**, in the superior node, already firing briskly at baseline, beats became faster and diastole filled with fluctuations. In the inferior node, sparse and subthreshold-dominated at baseline, the trace converted toward regular AP activity.

**Figure 8.**
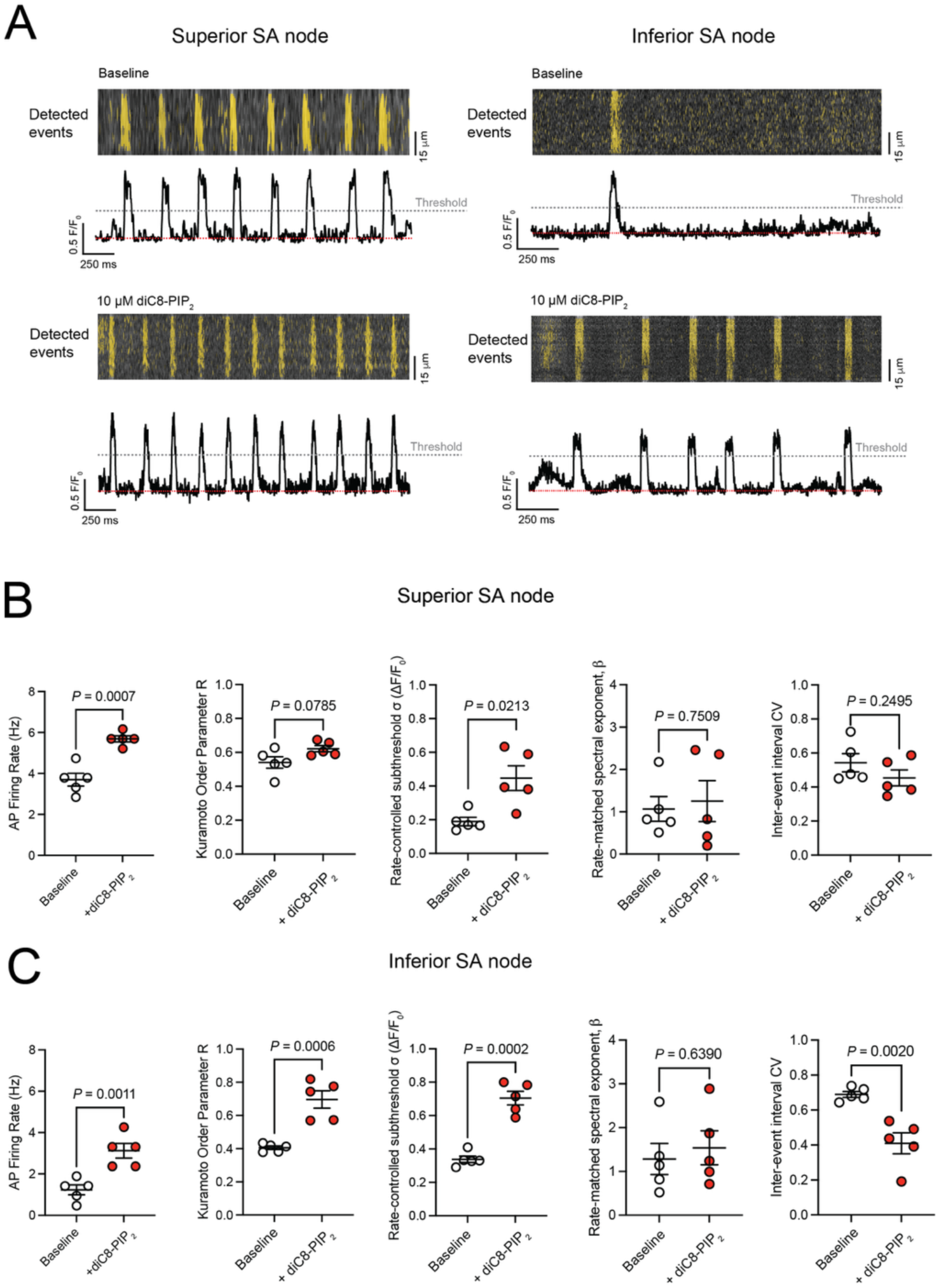
diC8-PIP_2_ accelerates firing and raises subthreshold noise in both poles but improves coherence only in the energy-poor inferior node. (A) Representative detected-event images and ΔF/F_0_ traces from the superior and inferior node at baseline and during 10 µM diC8-PIP_2_. (B) Superior node. Left to right: AP firing rate (*P* = 0.0007), Kuramoto order parameter R during diastole (*P* = 0.0785), mean rate-controlled subthreshold σ (*P* = 0.0213), rate-matched spectral exponent β (*P* = 0.7509), and inter-event interval CV (*P* = 0.2495) (*N* = 5). (C) Inferior node. Left to right: AP firing rate (*P* = 0.0011), R (*P* = 0.0006), σ (*P* = 0.0002), β (*P* = 0.6390), and inter-event interval CV (*P* = 0.0020) (*N* = 5). Each point represents one recording. Bars show means ± SEM. Exact *P*-values are shown. Each circle represents one animal; *N* denotes the number of mice.

The prediction held, but the effect was graded rather than simply present in one pole and absent in the other. In the superior node, AP firing rate rose modestly (*P* = 0.0007) and subthreshold σ roughly doubled, from 0.19 ± 0.02 to 0.45 ± 0.07 (*P* = 0.0213); yet diastolic coherence did not improve significantly (R = 0.54 ± 0.03 to R = 0.62 ± 0.02, *P* = 0.0785), and interval variability was unchanged (*P* = 0.2495) (**Figure 8B**). In the inferior node, firing rate roughly doubled (*P* = 0.0011), σ rose from 0.33 ± 0.02 to 0.70 ± 0.04 (*P* = 0.0002), coherence improved (R = 0.40 ± 0.01 to R = 0.69 ± 0.05, *P* = 0.0006), and interval variability fell (*P* = 0.0020) (**Figure 8C**). The spectral exponent β was unchanged in both poles (*P* = 0.7509 in superior; *P* = 0.6390 in inferior), so PIP_2_ raised the amplitude of the fluctuations without altering their character.

The difference between the poles is therefore not in whether the lipid acts, but what its action produces. Supplying PIP_2_ moved the noise coordinate in both poles, but that displacement was converted into a more ordered rhythm only in the pole beginning below its coherence optimum; coherence rose and interval variability fell together as they do at baseline. This is the same logic as the isoproterenol experiment, reached through a different molecular route: the effect of moving the noise coordinate depends on where the pole begins on the resonance curve.

Because diC8-PIP_2_ is supplied exogenously and bypasses PI4K entirely, the regional difference in response cannot directly reflect the ATP dependence of PIP_2_ synthesis; it instead reflects the baseline PIP_2_-occupation status of its target channels. PI4K, the rate-limiting enzyme for PIP_2_ synthesis, is ATP-dependent, with a half-maximal concentration (K_m_) of ∼209 µM (Tai *et al*., 2011). This places both poles above half-saturation: at roughly 400 µM, the inferior node runs at ∼66% of maximal velocity, and at roughly 800 µM the superior node runs at ∼79%. The poles therefore differ modestly in synthetic flux, but exhibit a greater difference in the sensitivity of that flux to ATP, since a given fractional change in ATP produces a 1.7-fold larger fractional change in synthesis inferiorly. Sustained over time, this lower and more ATP-sensitive synthetic rate would leave the inferior node with lower steady-state PIP_2_ and correspondingly reduced occupancy of its HCN channel PIP_2_-binding sites. That headroom, rather than any instantaneous difference in synthetic rate, is what the delivered lipid exploits. We did not measure PIP_2_ levels directly, but the inference accounts for exogenous PIP_2_ roughly doubling the firing rate inferiorly while yielding only a modest boost superiorly.

Taken together with **Figures 6** and **7**, the three experimental manipulations—isoproterenol, ivabradine, and PIP_2_—move the same two coordinates by different molecular routes and produce the outcomes each predicts. Isoproterenol raises noise, ivabradine lowers drive, and PIP_2_ relieves an energetic constraint on the conductance that carries the drive. In each case, the effect on coherence depends on where the pole began on the resonance curve, not simply on the direction of the intervention. Across all three interventions, β does not move, so the interventions displace the amplitude of the fluctuation without changing the process that generates it.

### PIP_2_ rescue of SA node firing requires I_f_

The preceding experiments establish that supplying PIP_2_ accelerates firing and raises subthreshold noise, most strongly in the ATP-poor inferior node. However, PIP_2_ facilitates HCN4 channel gating (Pian *et al*., 2006; Pian *et al*., 2007) and also modulates Ca_V_1.2, K_ATP,_ and NCX channels (Hilgemann & Ball, 1996), any of which could produce the same effect. If the rescue proceeds through I_f_, then blocking that current before supplying PIP_2_ should prevent it. We therefore imaged superior and inferior node preparations using a sequential, three-condition protocol: baseline, ivabradine, and ivabradine plus diC8-PIP_2_ (**Figure 9**).

**Figure 9.**
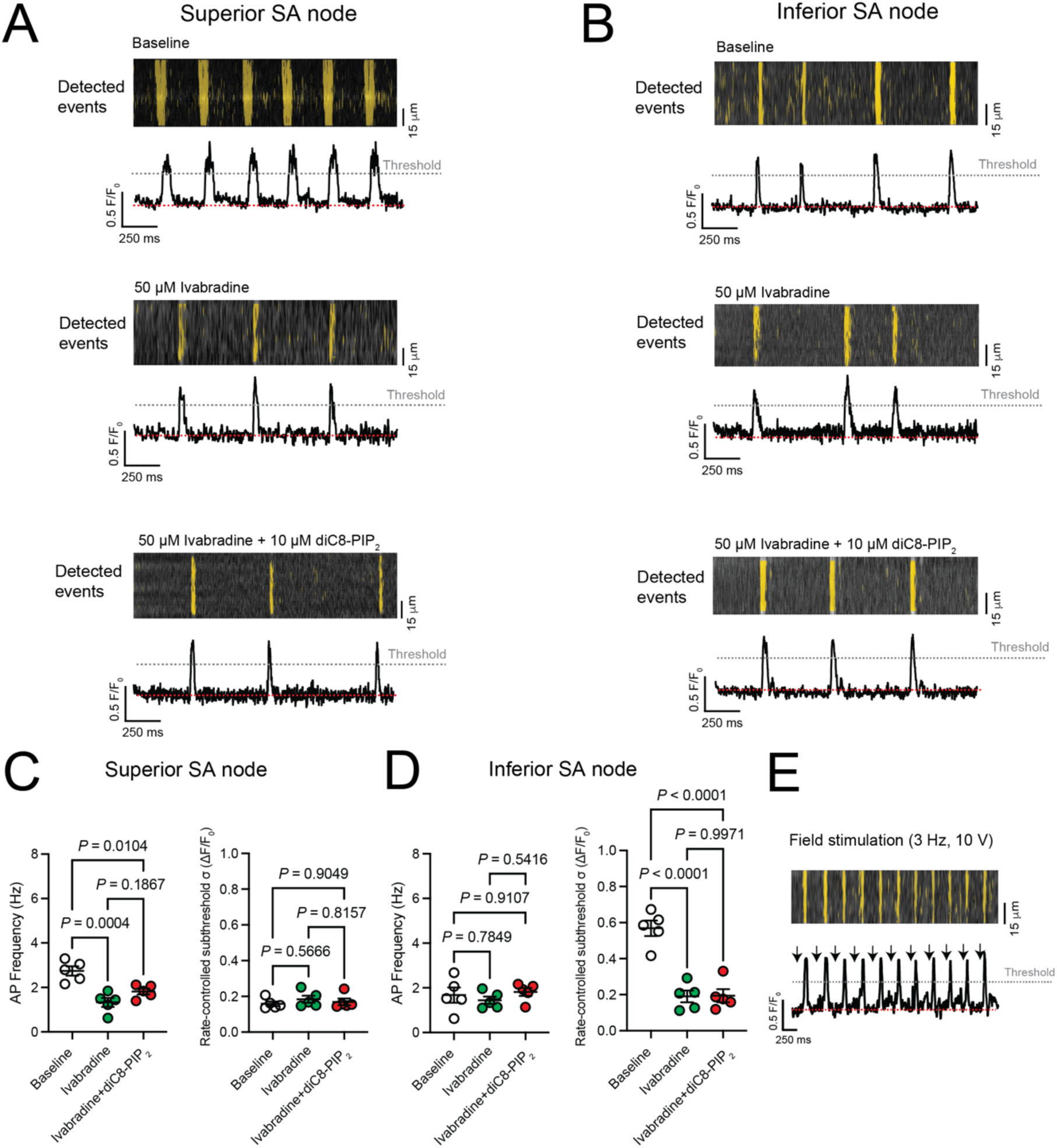
PIP_2_ acts on pacemaker firing through I_f_. (A, B) Representative detected-event images and corresponding ΔF/F_0_ traces from the superior SA node (A) and inferior SA node (B) under baseline conditions, during 50 µM ivabradine, and during 50 µM ivabradine + 10 µM diC8-PIP_2_. (C, D) AP firing frequency (left) and rate-controlled subthreshold σ (right) across the three conditions in the superior SA node (C) and inferior SA node (D) (*N* = 5 for both). (E) Evoked APs during 3 Hz, 10 V field stimulation in ivabradine-treated preparations. Each point represents one recording. Bars show means ± SEM. Exact *P*-values are shown. Each circle represents one animal; *N* denotes the number of mice.

Representative line scan images with detected events and the corresponding spatially averaged voltage records are shown for the superior (**Figure 9A**) and inferior (**Figure 9B**) nodes across all three conditions. In the superior node, ivabradine reduced AP frequency from 2.74 ± 0.20 to 1.33 ± 0.20 Hz (*P* = 0.0004) without changing subthreshold σ (*P* = 0.5666) (**Figure 9C**). Adding diC8-PIP_2_ in the presence of ivabradine did not restore AP frequency, which reached 1.81 ± 0.13 Hz and remained indistinguishable from ivabradine alone (*P* = 0.1867) at below baseline levels (*P* = 0.0104); subthreshold σ also remained unchanged (**Figure 9C**). In the inferior node, ivabradine did not significantly alter AP frequency (*P* = 0.7849) but markedly reduced subthreshold σ (*P* < 0.0001). Subsequent addition of diC8-PIP_2_ did not significantly alter AP frequency relative to ivabradine alone (*P* = 0.9107) and failed to restore subthreshold σ (*P* = 0.9971 vs. ivabradine), which remained significantly below baseline (*P* < 0.0001) (**Figure 9D**).

The failure to recover does not represent a loss of excitability. The same ivabradine-treated preparations captured evoked APs one-for-one under 3-Hz field stimulation (**Figure 9E**), so the cells remained capable of firing when driven; what was lost was the capacity to reach threshold on their own. These data support the hypothesis that PIP_2_ acts on pacemaker firing through I_f_: when the current is blocked, supplying PIP_2_ no longer restores the beat.

## Discussion

### Endogenous noise organizes rhythm in the intact node

The data presented in this study support a model in which the SA node is a spatially graded structure whose subthreshold fluctuations, generated within the tissue by its own conductances, serve as a functional variable rather than a residual problematic byproduct. Coherence among pacemaker cells is maximized at an intermediate amplitude of subthreshold voltage fluctuations. The superior and inferior SA node occupy distinct positions along this noise–coherence relationship, set by regional differences in bioenergetic support that also shape intrinsic excitability. The position on that relationship has two coordinates—the amplitude of the noise and the strength of the periodic drive—and interventions that move either one move coherence in the direction predicted by the resonance curve. This differs from the prevailing model of cardiac pacemaking, which treats the node as a deterministic oscillator network in which a dominant fast region entrains its neighbors and biological variability degrades performance. Instead, rhythm in the intact node is metabolically tuned and noise-assisted rather than purely entrained. The evidence for each element of this model, and its implications for pacemaker function, autonomic modulation and sinus node dysfunction, are considered below.

### Fluctuations are independent of the periodic drive

Stochastic resonance requires that fluctuations be independent of the weak periodic signal; if a single variable generated both, the non-monotonic coherence–σ relationship could instead be attributable to a nonlinearity in that variable rather than to a resonance between the two (Bezrukov & Vodyanoy, 1995; Clancy & Santana, 2020; Guarina *et al*., 2022; Pérez-Cervera *et al*., 2023; Kreider *et al*., 2025). Two considerations place Ca^2+^ cycling with the drive. The Ca^2+^ clock is the weak periodic oscillator carrying late diastolic depolarization toward threshold, so it structurally occupies the signal position. Mechanistically, it is not a passive ATP load but a self-reinforcing loop. SR Ca^2+^ release raises mitochondrial matrix Ca^2+^, activates matrix dehydrogenases, and contributes to the beat-locked ATP transients we have described in this tissue (Muñoz *et al*., 2026). These transients, in turn, provide the energy that ‘funds’ the SERCA activity and PI4K flux that the next release depends on. Cycle-locked positive feedback is the signature of a deterministic oscillator; noise sources do not possess feedback that entrain them to the cycle they perturb.

I_f_ occupies a different position, contributing to both noise amplitude and pacemaker drive. Its mean, deterministic component adds depolarizing current during diastole, while stochastic gating of the same channels generates the fluctuations riding on it. This dual role does not compromise independence: signal and noise can arise from the same molecular source, provided that the fluctuation is not a deterministic function of the periodic signal. Stochastic HCN gating carries no feedback that entrains it to the Ca^2+^ cycle, and the two components separate experimentally: blocking I_f_ predominantly impacted firing rate in the superior node without affecting subthreshold σ, and predominantly impacted σ in the inferior node, where the decrease in σ was the larger effect. The same current therefore acts predominantly as drive where the Ca^2+^ clock is strong and predominantly as a noise source where it is weak.

Our earlier work supports this directly, showing that thapsigargin abolished firing in every superior and inferior myocyte tested and eliminated the beat-locked ATP transients, yet left the frequency and amplitude of subthreshold fluctuations unchanged (Grainger *et al*., 2021; Muñoz *et al*., 2026). Spark-to-membrane coupling in these cells is weak but comparable between poles, yet sparks preceding an AP were more numerous in the superior node, the pole with the smaller subthreshold amplitude. Equal coupling with more sparks predicts larger superior σ; we observed the opposite. Eliminating SR release removes both the beat and the metabolic oscillation that funds it while leaving the fluctuation intact, positioning the noise upstream of the Ca^2+^ clock rather than alongside it (Muñoz *et al*., 2026). The Ca^2+^ clock is thus not only the periodic signal but the nonlinear amplifier that converts the fluctuation into a beat: because Ca^2+^-induced Ca^2+^ release (CICR) is regenerative, it acts as a threshold detector, converting a sufficiently large subthreshold event into a beat. Thapsigargin removes that amplifier and leaves its input intact. Independence is not unrestricted, since a clock that helps generate ATP also supports the conductances downstream of it, but that coupling acts over the minutes across which metabolic supply is set rather than within a beat. Signal and noise remain independent at the pacemaker frequency, and their slower coupling is precisely how perfusion positions a pole on the resonance curve.

Subthreshold noise is not attributable to a single conductance. Blocking I_f_ collapsed σ in the inferior node and produced a proportionally smaller effect on inferior node firing rate, while leaving superior node σ unchanged and halving its firing rate; thus, the ivabradine-sensitive conductance is necessary for the large subthreshold events of the inferior pole. Necessity is not identity. The single-channel conductance of HCN4 is small, and though correlated gating (Dekker & Yellen, 2006) generates variance above independent gating, it is an uncertain source of deflections of the amplitude resolved here. The interpretation more consistent with the data is that I_f_ supplies the initiating depolarization and that conductances activating at more depolarized potentials amplify it into the events we resolve. Coordinated opening of clustered Ca_V_1.3 channels in SA myocytes is a candidate mechanism for the discrete events, since correlated openings generate variance far greater than that of independent gating; coordinated multi-channel openings are resolved in these cells but lost with age (Vivas *et al*., 2026). According to this view, ivabradine abolishes the noise not by silencing its immediate source but by withdrawing the depolarization that recruits it. Ca_V_1.3-null mice exhibit bradycardia and sinus pauses, and loss-of-function mutations in the human Ca_V_1.3 channel subunit, CACNA1D, produce bradycardia (Mangoni *et al*., 2003; Baig *et al*., 2011), establishing a contribution of Ca_V_1.3 channels to reaching threshold. Whether they undergird the fluctuations is an open question, one that could be answered by measuring subthreshold events in Ca_V_1.3-null mice. Confirmation of spark suppression with ryanodine and an examination of voltage-clamp current-noise spectra at the maximum diastolic potential would discriminate the remaining candidates.

The regional contrast within the SA node has more to do with organization than amount. The superior node is not quiet; its largest individual events exceed those seen inferiorly. What differs is entrainment: fluctuations locked to the cycle join the deterministic trajectory and are removed by subtracting the trajectory (detrending), whereas unentrained fluctuations remain in the residual and contribute to σ. This distinction also reconciles our measurements with the local Ca^2+^ releases described in intact tissue by Okamura *et al*. (2026). Releases firing in register with the cycle are absorbed into the trajectory rather than the residual, so a spark-based noise substrate and the survival of σ under SERCA inhibition are not contradictory; they describe different components of the same signal. The Ca^2+^–mitochondrial loop suggests why this varies regionally, since closing this loop requires O_2_. Where supply is dense, ATP arrives in register with the beat and release stays phase-locked; where sparse, the loop is substrate-limited and release loses phase-lock. The thapsigargin results impose boundary conditions on what that decoupling can contribute: removing SR release altogether left σ unchanged in both poles, so the loss of phase-lock is better interpreted as a degradation of deterministic drive than as an increase in noise.

### σ reports how noise is expressed, not how much is generated

Voltage noise is current noise filtered by input impedance, so σ^2^ ≈ ⟨i^2^⟩·R_in_^2^: a cell with twice the input resistance shows twice the voltage noise for the same underlying stochastic current. I_f_ is active throughout diastole and acts as a shunt, lowering that resistance, so blocking it should raise R_in_ and inflate σ even with current noise unchanged. An interesting observation in this study is that ivabradine did not do this. σ was unchanged in the superior node and fell substantially in the inferior (**Figure 7B**). The expressed noise therefore did not follow input resistance. In the inferior node, the fall was large enough that the underlying current noise must itself have decreased when I_f_ was removed, rather than being an indication that the membrane merely stopped amplifying it. In the superior node, σ simply held, which is equally consistent with a modest change in R_in_ or with offsetting effects. The same logic applies regionally, with the inferior pole being more hyperpolarized with less I_f_.

Which conductances can carry fluctuations is constrained by arithmetic, and the targets are the subthreshold populations near 1.3 and 10.9 mV (Grainger *et al*., 2021). With ∼27 pF and diastolic input resistance of 0.5–1 GΩ (Vivas *et al*., 2026), the membrane time constant is ∼14–27 ms, so any current briefer than this is smoothed away before the membrane can fully respond to it. For HCN4, a unitary current of ∼0.04 pA at diastolic driving force across ∼10^4^ channels near *P_o_* = 0.1 yields ∼0.6 mV of independent gating noise; a mean pairwise correlation of only ρ = 0.001 inflates this threefold. However, because HCN4 gating is slow relative to the membrane, this appears as a beat-to-beat offset in the ramp rather than as resolvable events. For Ca_V_1.3, whose unitary current predicted by the Goldman–Hodgkin–Katz flux equation is ∼0.7 pA at the maximum diastolic potential, with up to seven coordinated openings resolved (Moreno *et al*., 2016; Vivas *et al*., 2026), four to seven sustained coordinated openings gives ∼1.4–2.5 mV, matching the smaller population, although brief Mode 1 openings of Ca_V_1.3 channels (0.5–1 ms) end long before the membrane has responded and deliver only 3–7% of that value. Correlated, sustained openings are therefore required rather than incidental. The larger events demand ∼11–22 pA, or 16–31 simultaneously open channels. Since SVFs survive SERCA inhibition undiminished (Grainger *et al*., 2021), this regeneration cannot be CICR, and is more likely voltage-dependent recruitment of additional Ca_V_1.3 channels by local depolarization. The implication of our findings is that amplification proceeds in two stages: a membrane-delimited subthreshold stage reaching ∼11 mV, and a SERCA-dependent stage in which the Ca^2+^ clock completes the beat. Of these, thapsigargin blocks only the second.

Computed over rate-matched bands, the spectral exponent β did not differ between poles and placed the subthreshold signal between pink (β = 1) and brown (β = 2) noise. Thus, β provides three constraints on the interpretation of the regional σ gradient.

*First*, the similar β values show that the σ gradient cannot be explained by the different beat rates of the two poles. The primary σ is measured in a fixed 100-ms window placed at the same phase of every cycle, so it cannot accumulate variance in proportion to diastolic duration and is immune to this bias by construction. The uncorrected measure in **Table 1** is not, and there the scaling can be bounded. For a correlated signal, variance accumulated over a finite window scales as T^(β−1)^, so σ scales as T^((β−1)/2)^; at β = 1.5 the exponent is 0.25. The inferior diastole is ∼3-fold longer than the superior, which would inflate an SD-based σ by about 30%, well short of the 2.6-fold regional difference measured in the paired comparison of **Table 1**. The σ gradient therefore cannot be attributed to the longer observation window imposed by the slower inferior rhythm.

*Second*, equal β constrains an explanation based primarily on membrane gain. If the larger inferior σ principally arose from higher inferior input resistance, the frequency above which the membrane starts to filter, 1/(2πR_in_C_m_), would differ between poles. That difference should alter the spectral exponent measured over a fixed, rate-matched frequency band. It does not. Thus, the regional σ difference includes a genuine difference in underlying current noise, rather than arising solely from differences in how the membrane converts current noise into voltage fluctuations.

*Third*, equal β does not imply equal fluctuation composition. Detected subthreshold events account for roughly 1/8^th^ of σ in the superior node but for a large majority of it in the inferior node, so the composition of the fluctuation is very nearly reversed between poles even though its spectral slope is unchanged. The two measures describe different aspects of the signal: the event partition reflects the fraction of total variance contributed by detected events across the full frequency range, whereas β is fitted only within the rate-matched band, which begins at twice the beat frequency.

This distinction explains why the striking regional difference in event composition does not produce a difference in β. A discrete event of duration τ has a Lorentzian spectrum with a corner frequency of 1/(2πτ). For events lasting tens of milliseconds, that corner falls at or below the lower edge of the fitted band, so most of the variance such events carry lies in the sub-corner plateau the β fit does not sample. Notching the beat fundamental and its first harmonics further removes more of this low-frequency region. The exponent is therefore estimated over a range in which the compositional difference is largely absent by construction, so its invariance is expected rather than paradoxical. β is therefore a proxy for neither σ nor composition; it reports a property of the fluctuation that remains constant across the poles, even as both amplitude and composition vary regionally.

### Coherence depends on both noise and depolarizing drive, and each pole sits at its own optimum

A pole’s position is set by two coordinates: unentrained fluctuation amplitude and drive strength. Coherence is a ridge over both, and a manipulation moves it according to which coordinate it displaces—predominantly noise for isoproterenol, downward drive for ivabradine, and the noise coordinate together with a probable drive component for PIP_2_. Viewed through this lens, the PIP_2_ result is a double dissociation: inferiorly, diC8-PIP_2_ raised σ and coherence together; superiorly it raised σ comparably but did not affect coherence. On a single noise axis, this is contradictory, since the inferior pole has the higher baseline σ and should therefore be the one pushed past its optimum. On a surface it is straightforward: the two poles begin at different positions, so the same displacement in σ carries the inferior node toward its optimum and the superior node away from its optimum. A drive contribution, plausible if PIP_2_ support of NCX strengthens the Ca^2+^ clock as discussed below, would move both poles along the second axis as well and would not be at odds with this two-coordinate interpretation. The poles therefore do not share one inverted-U relationship; their optima are region-specific, and each pole is best compared against its own inverted-U curve.

The flattening of the coherence–σ relationship under β-adrenergic drive bounds where this mechanism operates. Noise-dependent coordination appears to be a property of the node at rest, receding when catecholaminergic drive supplies coordination directly and the node tracks along the crest of the ridge rather than across it. The relevance of this regime differs across species. A mouse at 500–600 beats per minute spends little of its life there, whereas the human node at 60– 90 beats per minute operates in it continuously, with cycle lengths approaching an order of magnitude longer and correspondingly more diastole over which fluctuation can accumulate. The comparison is not a simple function of cycle length, since channel densities, current kinetics, and the number of coupled cells scale with body size, and a larger node averages over more of both; where a species sits on its own resonance spectrum is an empirical question. But the conditions that weaken drive relative to threshold, vagal slowing, sinus node disease, and age, extend that regime in any species.

Okamura *et al*. (2026) elegantly established the single-cell counterpart. Taking direct control of the fluctuation in isolated rabbit myocytes, they showed that injected white-noise currents awaken dormant cells and improve the rate and rhythm of slow, irregular cells, with the beneficial effect on regularity appearing at intermediate amplitude and lost at the extremes. Simultaneous membrane potential and Ca^2+^ recordings, supported by models spanning the Ca^2+^-release unit, the cell and the tissue, showed that the coupled-clock system amplifies subthreshold signaling under noise. In intact mouse tissue, they identified local Ca^2+^ releases as the noise substrate. Their basal-state data show the same opposition we found between poles: noise improves rhythmicity in slow-firing cells and degrades it in fast-firing cells.

We use ‘stochastic resonance’ in a bounded sense: coherence depends non-monotonically on unentrained fluctuation, the presence of a weak subthreshold periodic drive in the coupled membrane and Ca^2+^ clocks, and bidirectional movement of coherence in response to displacements along either coordinate. The alternative for an autonomous oscillator is coherence resonance, in which a noise optimum arises with no separable periodic input, and the inverted-U relationship alone does not distinguish the two. Neither does the two-coordinate dependence we report, since altering a parameter of an autonomous oscillator shifts its entire coherence–noise relationship and would produce the same signature. What would distinguish them is a periodic input whose frequency can be varied independently of the oscillator. Our preparation does not provide one: the two clocks are mutually entrained, so assigning one the role of signal and the other of detector is a modeling choice rather than an observation. Okamura *et al*. (2026) performed that experiment at the cellular level, showing that pacemaker myocytes follow an externally imposed periodic current over a bounded band of frequencies, which is what the bounded reading assumes. Classical mutual entrainment (Jalife, 1984; Anumonwo *et al*., 1991) also operates, and the two are complementary, since entrainment dissolves heterogeneity into a common rhythm whereas stochastic resonance requires it.

Our study and Okamura’s study are complementary along four axes: injected versus endogenous fluctuation, isolated cells versus intact tissue, Ca^2+^ versus membrane voltage, and cells sorted by firing behavior versus cells sorted by anatomical position. The first is the substantive one. An electrode establishes what a cell does when noise of known amplitude is supplied to it; it cannot establish whether the fluctuation a cell actually receives from its neighbors has the amplitude, spectrum, and spatial extent to do that work. This is the question a tissue measurement answers that an isolated cell cannot. The resonance spectra employed by Okamura *et al*. (2026) make the requirement explicit: for an endogenous fluctuation to act by the same route, its power must fall within the entrainable band. Our study satisfies this necessary condition, since β ≈ 1.5 places most of its power at the low frequencies Okamura *et al*. (2026) identify as effective. The last axis matters for a different reason. Sorting by firing behavior, as done by Okamura *et al*. (2026), establishes that the phenomenon exists; sorting by anatomical position, as we have done, asks what sets a cell’s place on the curve. The metabolic gradient we describe supplies an answer beyond the reach of behavioral sorting. That the agreement holds across species, indicator, and both sorting criteria makes it unlikely to be attributable to differences in methodology.

### Local energy supply positions each pole on its resonance curve through PIP_2_

Superior myocytes are better vascularized, more mitochondria-rich, and generate larger beat-locked ATP pulses (Muñoz *et al*., 2026). ATP funds deterministic drive through SERCA and the local-release ensemble, and it funds the PI4K flux that supplies the PIP_2_ necessary for HCN4 channel function, so a pole’s energy supply moves it along both coordinates of the resonance surface, with the dominant axis set by where it already sits relative to its own optimum. That PIP_2_ should report an energetic deficit early follows from kinetics. The Michaelis constant of PI4K for Mg·ATP is ∼209 µM (Tai *et al*., 2011), within the range of cytosolic ATP spanned by the node, so its activity varies across precisely the range over which the regional gradient operates. In contrast, Na^+^/K^+^-ATPase binds ATP at its high-affinity site in the micromolar range (Norby & Esmann, 1997) and remains saturated where PI4K does not. Membrane PIP_2_ therefore declines well before ionic gradients are threatened, and channels requiring it lose gating support while the cell remains, by the usual measures, energetically intact.

Because diC8-PIP_2_ bypasses PI4K, the regional selectivity of its effect follows from baseline occupancy rather than synthesis: a lower ATP ceiling leaves more unoccupied HCN4 sites for delivered lipid to fill, and raising σ improves coordination only in a pole that begins below its own optimum. Prior I_f_ block prevented diC8-PIP_2_ rescue (**Figure 9**), establishing that I_f_ is required, though not necessarily that it is the sole effector. PIP_2_ is not a selective HCN cofactor; it also regulates Ca_V_1.2, K_ATP_ and NCX (Hilgemann & Ball, 1996; Pian *et al*., 2006; Pian *et al*., 2007; Suh *et al*., 2012). Its impact on the K_ATP_ arm cannot explain the rescue, as PIP_2_ support of the resulting K^+^ efflux would hyperpolarize the maximum diastolic potential and slow firing—the opposite of what we observe. NCX is a different matter. PIP_2_ supports exchanger activity, and NCX carries the inward current generated by late diastolic SR Ca^2+^ release, so supplying PIP_2_ might strengthen the Ca^2+^ clock itself. Accordingly, PIP_2_ would raise drive as well as noise, moving a pole along both axes at once, as isoproterenol does.

Withdrawing energy supply tests the relationship from the other direction. A sub-pressor concentration of angiotensin II selectively reduces microvessel density in the superior node while sparing the inferior node, and produces bradycardia with an approximately 8-fold rise in interval variability (Manning *et al*., 2025). This intervention therefore reduces perfusion specifically in the pole with the lower subthreshold noise and the higher baseline coherence, and that pole becomes slower and more variable. This direction of change is consistent with the resonance model: reducing energy supply lowers deterministic drive, moving the superior pole away from its optimum and reducing coherence. By contrast, simply removing noise from a pole at or beyond its optimum should not degrade regularity.

### Tissue organization: the leading site, and what the inferior pole contributes

The regional difference in leading-site stability suggests an interpretation of the descending limb of the inverted-U curve that has no single-cell analogue. In tissue, excess fluctuation relative to drive should produce ectopic threshold crossings that compete with the propagating wave, so the failure mode reflects wandering initiation rather than beat-to-beat jitter, and leading-site stability becomes a readout of which limb—ascending or descending—a preparation occupies. This suggests a more economical reading of the node. We have written as though threshold, drive, and fluctuation reside within every myocyte, making the node a population of resonators whose performance we average. The regional data are equally consistent with the ingredients being anatomically segregated, with the superior pole supplying drive and threshold, and the inferior supplying unentrained fluctuation (delivered electrotonically through gap junctions), so that an inferior myocyte, which rarely reaches threshold, is not a failed oscillator but a current source. This possibility, which was raised when the regional gradient was first described (Grainger *et al*., 2021), remains untested. Because independent resonators predict the same gradient, our data are consistent with this possibility, but confirmation will require recording the two poles simultaneously in future work.

### Conclusions

The SA node is structurally heterogeneous and metabolically graded, and its coherence depends on the amplitude of its own subthreshold fluctuations. Perfusion sets a ceiling on ATP, ATP funds the deterministic drive that carries diastolic depolarization toward threshold, and the fluctuation that remains is not an unavoidable imperfection superimposed on the rhythm. It is generated by the tissue’s own conductances, and at an intermediate amplitude, it is a determinant of how reliably the heartbeat is timed. Three implications for the accepted models of cardiac pacemaking follow: 1) Variability in the node cannot be treated as an error term to be averaged away, since coherence depends on it non-monotonically. 2) A single dominant pacemaker region is not sufficient to describe initiation, since leading-site stability tracks position on the resonance curve rather than intrinsic rate. 3) The determinants of pacemaker rate and reliability extend beyond the membrane and Ca^2+^ clocks to the perfusion that supports them. The coupled-clock framework remains correct in what it describes; what these data add is that the output of this framework is graded by a bioenergetic supply chain.

## First author profile

Manuel F. Muñoz Camus, PhD, is a postdoctoral researcher in the Department of Physiology & Membrane Biology at the University of California, Davis, in the laboratory of Dr. Fernando Santana. He earned a BSc in Biological Sciences and a PhD in Physiological Sciences from the Pontificia Universidad Católica de Chile, where his doctoral research investigated the role of astrocytic Ca^2+^ waves in neurovascular coupling. During his postdoctoral training, Dr. Muñoz studied how S-nitrosylation of connexin-43 contributes to cardiac arrhythmogenesis and myocardial injury in Duchenne muscular dystrophy. In the Santana laboratory, his current research focuses on the sinoatrial node, where he investigates how regional ATP dynamics and bioenergetic gradients shape subthreshold electrical noise, ion-channel regulation, and coherence among pacemaker cells. Combining advanced optical imaging, genetically encoded biosensors, and electrophysiology, his work seeks to understand how cellular bioenergetics shapes cardiac excitability and pacemaking in health and disease.

## Data and Software Availability Statement

The GEVI LineScan Analyzer (v1.8.1) developed and used in this study is a single self-contained HTML file that runs locally on any modern web browser, with no installation, no server, and no data upload; all processing occurs client-side. Source code and subsequent versions are maintained at https://github.com/Santana-Lab-UC-Davis/GEVI-linescan-image-analyzer under the GNU General Public License v3.0. Its detection engine is a derived work of SparkMaster 2 (Tomek *et al*., 2023), reimplemented in JavaScript, extended to two symmetric polarity passes and retuned for voltage-indicator data; all analyses downstream of detection are original to this study. Version 1.8.1 is archived at Zenodo (https://doi.org/10.5281/zenodo.22085046). The archived release includes a seeded synthetic-data validation harness that reproduces the detection performance without access to the experimental recordings. All data generated or analyzed during this study are included in the manuscript.

## Competing interests

The authors declare that they have no competing interests.

## Author Contributions

- **Manuel F. Muñoz:** Conceptualization, investigation, formal analysis, software, visualization, writing.
- **Marc D. Binder:** Formal analysis, methodology, writing.
- **Christopher G. Wilson:** Formal analysis, methodology, writing.
- **L. Fernando Santana:** Conceptualization, supervision, funding acquisition, project administration, writing.

## Funding

National Heart, Lung, and Blood Institute (NHLBI) HL168874 grant to L.F.S.

## Acknowledgements

Portions of the source code for the custom analysis software were drafted with the assistance of a large language model (Claude Code, Anthropic). The authors are solely responsible for the study design, analysis, interpretation, and conclusions presented here. We thank Dr. Jody Martin (CVRI, UC Davis) for AAV production. We also thank Dr. William J. Spain, Dr. Declan Manning, and Dr. Paula Rhana for reading the manuscript.

---

**Supplementary Figure 1.**
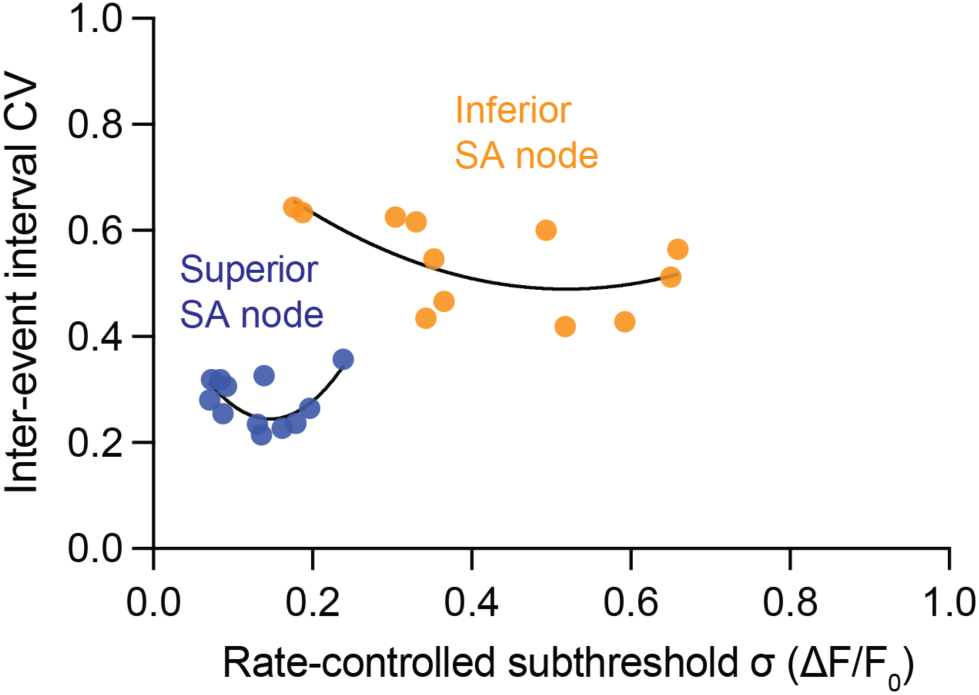
Interval variability traces the mirror image of diastolic coherence. Beat-to-beat interval variability (CV-IBI) plotted against subthreshold noise amplitude σ across recordings, with a quadratic fitted within each pole. Each pole traces a U with its minimum near the σ that maximizes diastolic coherence in Figure 4D, and CV-IBI is lower overall in the superior node.

## Notes

https://doi.org/10.5281/zenodo.22085046

### Competing Interest Statement

The authors have declared no competing interest.

### Summary of Updates

The manuscript has been reformatted for journal submission, including reordering of the Materials and Methods and Results sections. The description of ASAP5 baseline normalization has been clarified. A First Author Profile and Key Points Summary have been added, and the bioRxiv DOI has been included in the manuscript.

## References

Anumonwo JM, Delmar M, Vinet A, Michaels DC & Jalife J. (1991). Phase resetting and entrainment of pacemaker activity in single sinus nodal cells. Circ Res 68, 1138–1153.

Baig SM, Koschak A, Lieb A, Gebhart M, Dafinger C, Nurnberg G, Ali A, Ahmad I, Sinnegger-Brauns MJ, Brandt N, Engel J, Mangoni ME, Farooq M, Khan HU, Nurnberg P, Striessnig J & Bolz HJ. (2011). Loss of Ca(v)1.3 (CACNA1D) function in a human channelopathy with bradycardia and congenital deafness. Nat Neurosci 14, 77–84.

Bezrukov SM & Vodyanoy I. (1995). Noise-induced enhancement of signal transduction across voltage-dependent ion channels. Nature 378, 362–364.

Breakspear M, Heitmann S & Daffertshofer A. (2010). Generative models of cortical oscillations: neurobiological implications of the kuramoto model. Front Hum Neurosci 4, 190.

Brennan JA, Chen Q, Gams A, Dyavanapalli J, Mendelowitz D, Peng W & Efimov IR. (2020). Evidence of Superior and Inferior Sinoatrial Nodes in the Mammalian Heart. JACC Clin Electrophysiol 6, 1827–1840.

Bucchi A, Baruscotti M & DiFrancesco D. (2002). Current-dependent block of rabbit sino-atrial node I(f) channels by ivabradine. J Gen Physiol 120, 1–13.

Bucchi A, Tognati A, Milanesi R, Baruscotti M & DiFrancesco D. (2006). Properties of ivabradine-induced block of HCN1 and HCN4 pacemaker channels. J Physiol 572, 335– 346.

Bychkov R, Juhaszova M, Calvo-Rubio Barrera M, Donald LAH, Coletta C, Shumaker C, Moorman K, Sirenko ST, Maltsev AV, Sollott SJ & Lakatta EG. (2022). The Heart’s Pacemaker Mimics Brain Cytoarchitecture and Function: Novel Interstitial Cells Expose Complexity of the SAN. JACC Clin Electrophysiol 8, 1191–1215.

Bychkov R, Juhaszova M, Tsutsui K, Coletta C, Stern MD, Maltsev VA & Lakatta EG. (2020). Synchronized Cardiac Impulses Emerge From Heterogeneous Local Calcium Signals Within and Among Cells of Pacemaker Tissue. JACC Clin Electrophysiol 6, 907–931.

Clancy CE & Santana LF. (2020). Evolving Discovery of the Origin of the Heartbeat: A New Perspective on Sinus Rhythm. JACC Clin Electrophysiol 6, 932–934.

Csepe TA, Zhao J, Hansen BJ, Li N, Sul LV, Lim P, Wang Y, Simonetti OP, Kilic A, Mohler PJ, Janssen PM & Fedorov VV. (2016). Human sinoatrial node structure: 3D microanatomy of sinoatrial conduction pathways. Prog Biophys Mol Biol 120, 164–178.

Dekker JP & Yellen G. (2006). Cooperative gating between single HCN pacemaker channels. J Gen Physiol 128, 561–567.

DiFrancesco D. (1986). Characterization of single pacemaker channels in cardiac sino-atrial node cells. Nature 324, 470–473.

Dobrzynski H, Li J, Tellez J, Greener ID, Nikolski VP, Wright SE, Parson SH, Jones SA, Lancaster MK, Yamamoto M, Honjo H, Takagishi Y, Kodama I, Efimov IR, Billeter R & Boyett MR. (2005). Computer three-dimensional reconstruction of the sinoatrial node. Circulation 111, 846–854.

Efimov IR, Fedorov VV, Joung B & Lin SF. (2010). Mapping cardiac pacemaker circuits: methodological puzzles of the sinoatrial node optical mapping. Circ Res 106, 255–271.

Fedorov VV, Glukhov AV, Chang R, Kostecki G, Aferol H, Hucker WJ, Wuskell JP, Loew LM, Schuessler RB, Moazami N & Efimov IR. (2010). Optical mapping of the isolated coronary-perfused human sinus node. J Am Coll Cardiol 56, 1386–1394.

Fedorov VV, Schuessler RB, Hemphill M, Ambrosi CM, Chang R, Voloshina AS, Brown K, Hucker WJ & Efimov IR. (2009). Structural and functional evidence for discrete exit pathways that connect the canine sinoatrial node and atria. Circ Res 104, 915–923.

Fenske S, Hennis K, Rotzer RD, Brox VF, Becirovic E, Scharr A, Gruner C, Ziegler T, Mehlfeld V, Brennan J, Efimov IR, Pauza AG, Moser M, Wotjak CT, Kupatt C, Gonner R, Zhang R, Zhang H, Zong X, Biel M & Wahl-Schott C. (2020). cAMP-dependent regulation of HCN4 controls the tonic entrainment process in sinoatrial node pacemaker cells. Nat Commun 11, 5555.

Ferreiro M, Petrosky AD & Escobar AL. (2012). Intracellular Ca2+ release underlies the development of phase 2 in mouse ventricular action potentials. Am J Physiol Heart Circ Physiol 302, H1160–1172.

Glukhov AV, Fedorov VV, Anderson ME, Mohler PJ & Efimov IR. (2010). Functional anatomy of the murine sinus node: high-resolution optical mapping of ankyrin-B heterozygous mice. Am J Physiol Heart Circ Physiol 299, H482–491.

Grainger N, Guarina L, Cudmore RH & Santana LF. (2021). The Organization of the Sinoatrial Node Microvasculature Varies Regionally to Match Local Myocyte Excitability. Function (Oxf) 2, zqab031.

Guarina L, Moghbel AN, Pourhosseinzadeh MS, Cudmore RH, Sato D, Clancy CE & Santana LF. (2022). Biological noise is a key determinant of the reproducibility and adaptability of cardiac pacemaking and EC coupling. J Gen Physiol 154.

Hao YA, Lee S, Roth RH, Natale S, Gomez L, Taxidis J, O’Neill PS, Villette V, Bradley J, Wang Z, Jiang D, Zhang G, Sheng M, Lu D, Boyden E, Delvendahl I, Golshani P, Wernig M, Feldman DE, Ji N, Ding J, Sudhof TC, Clandinin TR & Lin MZ. (2024). A fast and responsive voltage indicator with enhanced sensitivity for unitary synaptic events. Neuron 112, 3680–3696 e3688.

Hilgemann DW & Ball R. (1996). Regulation of cardiac Na+,Ca2+ exchange and KATP potassium channels by PIP2. Science 273, 956–959.

Honjo H, Boyett MR, Kodama I & Toyama J. (1996). Correlation between electrical activity and the size of rabbit sino-atrial node cells. J Physiol 496 (Pt 3), 795–808.

Jalife J. (1984). Mutual entrainment and electrical coupling as mechanisms for synchronous firing of rabbit sino-atrial pace-maker cells. J Physiol 356, 221–243.

Kim MS, Maltsev AV, Monfredi O, Maltseva LA, Wirth A, Florio MC, Tsutsui K, Riordon DR, Parsons SP, Tagirova S, Ziman BD, Stern MD, Lakatta EG & Maltsev VA. (2018). Heterogeneity of calcium clock functions in dormant, dysrhythmically and rhythmically firing single pacemaker cells isolated from SA node. Cell Calcium 74, 168–179.

Kreider M, Lindner B & Thomas PJ. (2025). Q-functions, synchronization, and Arnold tongues for coupled stochastic oscillators. Chaos 35.

Lakatta EG, Maltsev VA & Vinogradova TM. (2010). A coupled system of intracellular Ca^2+^ clocks and surface membrane voltage clocks controls the timekeeping mechanism of the heart’s pacemaker. Circ Res 106, 659–673.

Li N, Hansen BJ, Csepe TA, Zhao J, Ignozzi AJ, Sul LV, Zakharkin SO, Kalyanasundaram A, Davis JP, Biesiadecki BJ, Kilic A, Janssen PML, Mohler PJ, Weiss R, Hummel JD & Fedorov VV. (2017). Redundant and diverse intranodal pacemakers and conduction pathways protect the human sinoatrial node from failure. Sci Transl Med 9.

Linscheid N, Logantha S, Poulsen PC, Zhang S, Schrolkamp M, Egerod KL, Thompson JJ, Kitmitto A, Galli G, Humphries MJ, Zhang H, Pers TH, Olsen JV, Boyett M & Lundby A. (2019). Quantitative proteomics and single-nucleus transcriptomics of the sinus node elucidates the foundation of cardiac pacemaking. Nat Commun 10, 2889.

Maltsev VA & Stern MD. (2022). The paradigm shift: Heartbeat initiation without “the pacemaker cell”. Front Physiol 13, 1090162.

Mangoni ME, Couette B, Bourinet E, Platzer J, Reimer D, Striessnig J & Nargeot J. (2003). Functional role of L-type Cav1.3 Ca2+ channels in cardiac pacemaker activity. Proc Natl Acad Sci U S A 100, 5543–5548.

Manning D, Rivera EJ, Rhana P, Matsumoto C, Fong Z, Thai PN, Munoz MF, Contreras JE, Kim S, Grainger N, Chiamvimonvat N, Bautista GM & Santana LF. (2025). Microvascular Rarefaction in the Sinoatrial Node: A Potential Mechanism for Pacemaker Dysfunction in Early HFpEF. JACC Clin Electrophysiol 11, 2107–2133.

Moreno CM, Dixon RE, Tajada S, Yuan C, Opitz-Araya X, Binder MD & Santana LF. (2016). Ca(2+) entry into neurons is facilitated by cooperative gating of clustered CaV1.3 channels. Elife 5.

Muñoz MF, Matsumoto C, Rhana P, Manning D, Bautista GM, Collier DM & Santana LF. (2026). Beat-locked ATP microdomains in the sinoatrial node map a Ca2+-timed energetic hierarchy and regional pacemaker roles. J Gen Physiol 158.

Norby JG & Esmann M. (1997). The effect of ionic strength and specific anions on substrate binding and hydrolytic activities of Na,K-ATPase. J Gen Physiol 109, 555–570.

Okamura A, He IK, Maltsev AV, Bychkov R, Tagirova S, Wang M, Maltsev AV, Stern MD, Lakatta EG & Maltsev VA. (2026). Cardiac Pacemaker Cells Harness Stochastic Resonance to Avoid Sinus Arrest. Circ Res 139, e328905.

Pérez-Cervera A, Gutkin B, Thomas PJ & Lindner B. (2023). A universal description of stochastic oscillators. Proc Natl Acad Sci U S A 120, e2303222120.

Pian P, Bucchi A, Decostanzo A, Robinson RB & Siegelbaum SA. (2007). Modulation of cyclic nucleotide-regulated HCN channels by PIP(2) and receptors coupled to phospholipase C. Pflugers Arch 455, 125–145.

Pian P, Bucchi A, Robinson RB & Siegelbaum SA. (2006). Regulation of gating and rundown of HCN hyperpolarization-activated channels by exogenous and endogenous PIP2. J Gen Physiol 128, 593–604.

Pikovsky AS & Kurths J. (1997). Coherence Resonance in a Noise-Driven Excitable System. Physical Review Letters 78, 775–778.

Reddy GR, Ren L, Thai PN, Caldwell JL, Zaccolo M, Bossuyt J, Ripplinger CM, Xiang YK, Nieves-Cintron M, Chiamvimonvat N & Navedo MF. (2022). Deciphering cellular signals in adult mouse sinoatrial node cells. iScience 25, 103693.

Rhana P, Matsumoto C & Santana LF. (2026). Demonstration of beat-to-beat, on-demand ATP synthesis in ventricular myocytes reveals sex-specific mitochondrial and cytosolic dynamics. J Physiol.

Santana LF & Earley S. (2026). Energetic microdomains and the vascular control of neuronal and muscle excitability: Toward a unified model. J Physiol.

Suh BC, Kim DI, Falkenburger BH & Hille B. (2012). Membrane-localized beta-subunits alter the PIP2 regulation of high-voltage activated Ca2+ channels. Proc Natl Acad Sci U S A 109, 3161–3166.

Swift LM, Kay MW, Ripplinger CM & Posnack NG. (2021). Stop the beat to see the rhythm: excitation-contraction uncoupling in cardiac research. Am J Physiol Heart Circ Physiol 321, H1005–H1013.

Tai AW, Bojjireddy N & Balla T. (2011). A homogeneous and nonisotopic assay for phosphatidylinositol 4-kinases. Anal Biochem 417, 97–102.

Telgkamp P, Cao YQ, Basbaum AI & Ramirez JM. (2002). Long-term deprivation of substance P in PPT-A mutant mice alters the anoxic response of the isolated respiratory network. J Neurophysiol 88, 206–213.

Tellez JO, Dobrzynski H, Greener ID, Graham GM, Laing E, Honjo H, Hubbard SJ, Boyett MR & Billeter R. (2006). Differential expression of ion channel transcripts in atrial muscle and sinoatrial node in rabbit. Circ Res 99, 1384–1393.

Tomek J, Nieves-Cintron M, Navedo MF, Ko CY & Bers DM. (2023). SparkMaster 2: A New Software for Automatic Analysis of Calcium Spark Data. Circ Res 133, 450–462.

Vivas O, Baudot M, Madden R, Pinon-Teal WL, Hunt MS, Choi S, Pournejati R, Flores-Tamez VA, Santana LF & Moreno CM. (2026). Aging Disrupts L-type Ca(2+) Channel Organization and Function in Pacemaker Cells. Circ Res 139, e327894.

Zanella S, Doi A, Garcia AJ, 3rd, Elsen F, Kirsch S, Wei AD & Ramirez JM. (2014). When norepinephrine becomes a driver of breathing irregularities: how intermittent hypoxia fundamentally alters the modulatory response of the respiratory network. J Neurosci 34, 36–50.

